# Global organisation of structural covariance networks derived from parcellated cortical surface area in atypical populations

**DOI:** 10.1101/2025.10.29.685428

**Authors:** William Roseby, Chris Racey, Jamie Ward

**Affiliations:** School of Psychology, University of Sussex

## Abstract

Higher-order features of brain organisation are powerful measures for understanding the relationship between brain and experience. In particular, the global arrangement of structural features of the cortex provides insight into neurodevelopmental processes that underlie individual differences in perception and cognition. Structural covariance networks (SCNs), which capture regional coordination of brain morphometry, are an efficient method to derive global properties of the cortex. However, their interpretation relies on an array of methodological choices that are often inconsistent between studies. Using a hierarchically-clustered version of the Human Connectome Project (HCP) atlas, we constructed SCNs of regional cortical surface area for groups with four different conditions – synaesthesia, autism, early psychosis, and anxiety or depression – and compared global network properties with those of age- and sex-matched controls. SCNs for synaesthesia and autism showed globally stronger connectivity, with specific increases at moderate cortical distances, as well as lower network complexity. Conversely, the SCN for early psychosis showed a globally lower connectivity and a greater complexity, while depression and anxiety showed few differences compared to controls. The results for autism and depression were replicated across two datasets each. These findings support the notion that synaesthesia and autism share neurodevelopmental mechanisms, while psychosis may involve a diverging process. This study is also an important proof of principle for analysing diverse populations under one methodological framework.

## 1 Introduction

A driving question in cognitive neuroscience is how the organisation of the brain is related to differences in perception, cognition, emotion, and overall experience. One method of characterising brain organisation is to examine the properties of cortical networks, where nodes represent specific regions of cortex and connections reflect the degree of association between them. This data-driven approach contrasts the examination of specific regions in isolation, which can improve statistical power but also introduce bias and constrain the hypothesis space (Wilson et al., 2024). The construction of global networks serves as a form of dimensionality reduction, rendering interpretable the higher-order properties of the brain which may be more sensitive to perturbation and more important for high-level cognitive processes (Montembeault et al., 2012; King et al., 2021; Vetter et al., 2025). Therefore, they may be particularly primed to uncover neural organisational principles that are strongly related to experiential qualities (Mišić and Sporns, 2016; Tompson et al., 2018).

Structural covariance networks (SCNs) are networks computed using structural MRI data by deriving the correlations between the morphometrics of different brain regions. Being based on structural features like grey matter volume, cortical thickness, or cortical surface area, these correlations capture the degree of coordinated growth of different regions that is driven by neurodevelopmental processes (Mechelli et al., 2005; Zielinski et al., 2010; Alexander-Bloch, Giedd and Bullmore, 2013). However, this structural association has also been shown to reflect genetic influences (Schmitt et al., 2008), experience-dependent plasticity and aging (Draganski et al., 2004; Montembeault et al., 2012), as well as white matter connectivity (Gong et al., 2012; Moura et al., 2017) and functional connectivity (Bhojraj et al., 2010; Alexander-Bloch, Raznahan, Bullmore and Giedd, 2013; Geerligs et al., 2016). Therefore, structural covariance presents a general yet powerful measure for cortical association and connectivity. Indeed, SCN topology has been used successfully to predict the status of various conditions like Alzheimer’s, Parkinson’s and schizophrenia (Diao et al., 2024; Mongay-Ochoa et al., 2025; Vetter et al., 2025). This power of this approach is augmented by its methodological advantages, which include the wider availability, lower computational cost, and greater signal-to-noise ratio of structural MRI compared to functional and diffusion MRI (Bethlehem et al., 2017; Diao et al., 2024). However, SCN studies on some conditions like autism have shown conflicting results (Bethlehem et al., 2017; Cai et al., 2021; Wang et al., 2022), which is likely attributable to methodological choices such as the structural measure, cortical parcellation scheme, correlation metric, and the application of thresholding, binarisation, or selection of positive or negative correlations (Liang et al., 2009; Zalesky et al., 2010; Joshi et al., 2010; Gong et al., 2012; Geerligs et al., 2016; Carmon et al., 2020). Therefore, there is a need to examine a variety of phenotypes following the same analysis pipeline.

The application of a consistent methodological framework across various cognitive phenotypes also bears important conceptual implications. Atypical cognitive phenotypes, such as autism, synaesthesia, psychosis, and depression, can serve as models to understand the relationship between brain organisation and experience. Yet, addressing how these diverse phenotypes are related through neurodevelopmental pathways may inherently require a comparative approach (Ward, 2021). Bayesian brain and predictive processing accounts have offered a unifying conceptual framework for each of these conditions, with changes to the relative balance of prior predictions and sensory information proposed to underlie an assortment of cognitive features (Adams et al., 2013; Lawson et al., 2014; Seth, 2014; Van de Cruys et al., 2017; Sterzer et al., 2018; Friston, 2020). Importantly, some have proposed a diametrical model of autism and psychosis, positing they lie at opposite ends of a single continuum (Crespi and Badcock, 2008; Rządeczka et al., 2023). However, how these theoretical alterations to inference relate to brain organisation is unclear, limiting the strength of conclusions regarding their neurodevelopmental overlap. Yet, some empirical evidence already reveals patterns between these atypical experiences. For example, autistic populations have higher rates of synaesthesia (Baron-Cohen et al., 2013; Hughes et al., 2017), a phenomenon where a perception of one kind induces a percept in a different sensory modality. Those with synaesthesia also report higher levels of autistic traits (Ward et al., 2017; van Leeuwen et al., 2019), and there is evidence for a genetic association of synaesthesia with autism, but not with schizophrenia (Nugent and Ward, 2022). Interestingly, autism and psychotic disorders also co-occur at an elevated rate, but whether this reflects troublesome diagnosis is unclear (Zhou et al., 2019; Trevisan et al., 2020), and evidence for a genetic overlap is inconclusive (Chandrasekhar et al., 2020; Jutla et al., 2022). Neurobiological findings of cortical networks tend to diverge, with autism showing greater between-network and short-range connectivity particularly involving the default mode network (DMN) (Müller et al., 2011; Hong et al., 2019; Ilioska et al., 2023; Weber et al., 2024; Lee et al., 2025), and psychosis / schizophrenia showing a pattern of increased connectivity within and between the DMN, salience network (SN) and fronto-parietal network (FPN) (Whitfield-Gabrieli et al., 2009; Hare et al., 2019; Del Fabro et al., 2021; Menon et al., 2023). A consistent finding is that autism, synaesthesia and psychosis are all associated with elevated rates of depression and anxiety (Achim et al., 2011; Hollocks et al., 2019; Li et al., 2020; Neufeld et al., 2025). However, while the former are all highly heritable (Bosley and Eagleman, 2015; Tick et al., 2016; Hilker et al., 2018), depression and anxiety are shaped more strongly by life circumstances, making it difficult to disentangle neurodevelopmental influences from societal and environmental factors like stigmatisation. A comparative approach may also be uniquely beneficial in this respect.

In this study, we used SCNs to examine the global structural organisation of the brain across a variety of atypical conditions including synaesthesia, autism, early psychosis, depression and anxiety, which were individually compared with age- and sex-matched controls. SCNs were constructed using cortical surface surface area from the Human Connectome Project (HCP) atlas parcellation, which was first clustered to generate a smaller set of 62 contiguous cortical regions. Edge weights in the SCNs were then calculated as the group-wise partial correlations between surface area values. Surface area was used because it has been shown to be substantially more robust for SCN pipelines than other measures (Carmon et al., 2020). Parcellated surface area is also less studied than cortical thickness, but these features may have distinct genetic influences (Panizzon et al., 2009; van der Meer et al., 2020) and surface area likely better reflect the group differences seen in synaesthesia and autism (Hazlett et al., 2017; Ward et al., 2024). Similarly, we used partial correlations over standard pearson correlations because they better represent pairwise relationships and have shown to align more closely with structural connectivity (Wheland et al., 2012; Masuda et al., 2018). In addition to comparing abstract network topology, the role of physical organisation was also analysed by aligning network weights with cortical distances. In agreement with some prior studies on cortical thickness (Hänggi et al., 2011; Wang et al., 2022), we found increased structural correlation in surface area for synaesthesia and autism groups compared to controls. Contrastingly, there was evidence for globally decreased structural correlation in early psychosis, and no group differences in anxiety and depression samples, supporting the notion that structural correlations more strongly reflect neurodevelopmental processes. The results were consistent across several datasets where they were available, suggesting robustness of the analysis.

## 2 Methods

The study methods were pre-registered on OSF following preliminary findings with the Syn100 dataset (https://osf.io/xzwav/overview). In addition to examining the present atypical populations, we originally planned to incorporate network comparisons across ages and between sexes. For brevity we exclude them here. We also restricted our analysis to surface area rather than cortical thickness. We do not include our analysis on the local clustering coefficient as it provided little additional insight over the global measure. Otherwise, our analyses conform to the pre-registrated methods. We expanded on network complexity as one of our stated exploratory analyses.

### 2.1 Data collection

No primary data collection occurred in this study. Seven specific datasets were accessed for secondary analysis. These included: synaesthesia 100 brains (Syn100) (Racey et al., 2023), the Autism Brain Imaging Data Exchange (ABIDE)-I (Di Martino et al., 2014), ABIDE-II (Di Martino et al., 2017), the Human Connectome Project for Early Psychosis (HCP-EP) (Prunier and Shenton Martha; Breier, n.d.), the Boston Adolescent Neuroimaging of Depression and Anxiety (BANDA) (Hubbard et al., 2024), Dimensional Connectomics of Anxious Misery (DCAM), and Perturbation of the Treatment of Resistant Depression Connectome by Fast-Acting Therapies (PDC). Some data were also acquired from the HCP Development/Aging (HCP-DA) (Harms et al., 2018) and HCP Young Adult (HCP-YA) (Van Essen et al., 2013) cohorts for supplementing numbers of healthy controls.

The Syn100 dataset was already available to us from a previous study (Racey et al., 2023). ABIDE-I and ABIDE-II data were downloaded from the Neuroimaging Tools & Resources Collaboratory (NITRC) amazon web service (aws) S3 bucket, located at s3://fcp-indi/data/Projects/ABIDE and s3://fcp-indi/data/Projects/ABIDE2 respectively. HCP-EP, BANDA, DCAM, PDC, and HCP-D were downloaded from the NIMH data archive (NDA). HCP-YA data were downloaded from the ConnectomeDB aws S3 bucket, located at s3://hcp-openaccess/HCP1200/. Collected data included T1w MRI scans in.nii.gz format, HCP pre-processed structural data, and phenotypic data including age, sex, and condition status. In the case of PDC, only pre-treatment data were collected.

### 2.2 Statement of ethics

As stated in the original study (Ward et al., 2024), participants in the Syn100 dataset gave informed consent for both participation and data sharing and the project was approved by the Brighton and Sussex Medical School (BSMS) Research Governance and Ethics Committee. Data from the NDA was accessed for research purposes following the Data Use Certification with approval from the NDA Data Access Committee. Data from the Neuroimaging Tools & Resources Collaboratory (NITRC) was accessed following registration. See Acknowledgements for further details on accessing each dataset.

### 2.3 Participant selection

The sample size for the smallest dataset was *N* = 102 participants in the synaesthete group. In order to facilitate comparable analyses, we constrained every dataset to this same sample size. Some additional criteria were placed on certain datasets: 18+ years of age in ABIDE, and for the BANDA participants with anxiety and depression were placed into a single group termed Mood Disorder (MD).

Participants were selected via a 1:1 optimal matching procedure by sex and age in most cases. The ‘affected’ group (with a positive diagnosis or assessment) were first selected by matching to the synaesthete sample; BANDA was matched according to sex only. The control group were then selected via matching the affected group of *N* = 102 to available control participants. The control pool for the Syn100 dataset was the same as the original study, which was *N* = 650 participants comprising a mixture of HCP-YA and HCP-DA subjects, as well as some participants scanned at the University of Sussex (Ward et al., 2024). For HCP-EP, BANDA, DCAM and PDC, there were insufficient internal control participants for a 1:1 matching. In these cases, all internal controls were used and the remaining affected group were matched to HCP-YA and HCP-DA participants. Participants were never used twice across datasets. This matching procedure produced groups of *N* = 102 for all datasets, with very close age and sex distributions for each paired control and affected groups. There was not more than a year between mean ages, and < 7% difference in sex proportions in the worst case (Table 1).

**Table 1:** Demographics and data sources for control and affected groups for each dataset. UoS = University of Sussex.

| Sample | Age (mean years +/- SD) | Sex (% female) | Data sources |
| --- | --- | --- | --- |
| Syn100 | Control: 35.34 (+/-13.15)<br>Aff: 35.38 (+/-13.20) | Control: 78.43<br>Aff: 77.45 | Control: UoS, HCP-YA, HCP-DA<br>Aff: UoS |
| ABIDE-I | Control: 29.95 (+/-7.14)<br>Aff: 30.82 (+/-8.73) | Control: 17.65<br>Aff: 16.67 | Control: ABIDE-I<br>Aff: ABIDE-I |
| ABIDE-II | Control: 28.65 (+/-11.17)<br>Aff: 28.37 (+/-11.44) | Control: 13.73<br>Aff: 13.73 | Control: ABIDE-II<br>Aff: ABIDE-II |
| HCP-EP | Control: 23.28 (+/-3.92)<br>Aff: 23.40 (+/-3.59) | Control: 53.92<br>Aff: 46.08 | Control: HCP-EP, HCP-YA, HCP-DA<br>Aff: HCP-EP |
| BANDA | Control: 15.43 (+/-1.07)<br>Aff: 15.66 (+/-0.87) | Control: 70.59<br>Aff: 77.45 | Control: BAND A, HCP-DA<br>Aff: BAND A |
| DCAM | Control: 30.54 (+/-9.67)<br>Aff: 30.82 (+/-8.88) | Control: 75.49<br>Aff: 76.47 | Control: DCAM, HCP-YA, HCP-DA<br>Aff: DCAM |
| PDC | Control: 34.10 (+/-10.72)<br>Aff: 36.59 (+/-12.73) | Control: 74.51<br>Aff: 77.45 | Control: PDC, HCP-YA, HCP-DA<br>Aff: PDC |

A power analysis for comparison of *n* = 1891 network correlations suggested that a *β* = 0.8 (*α* =.05) would be sufficient to detect a difference in means of *d* = 0.091 and a difference in variances of *f* = 0.046. A simplified version of a permutation test suggested that for a comparison of means from two sets of *n* = 102 observations, each drawn randomly from a normal distribution, the estimated power is approximately 0.8 (*α* =.05) for a difference in distribution means of *d* = 0.4. Therefore, our analysis appeared moderately well-powered to detect medium effect sizes, although this could differ for derived measures that do not follow a normal distribution.

### 2.4 MRI preprocessing and feature extraction

Preprocessing of structural data differed depending on whether the data were raw T1w scans (ABIDE-I, ABIDE-II, HCP-EP, DCAM) or were already preprocessed with the HCP pipeline (Syn100, BANDA, PDC). The HCP preprocessing pipeline is a multi-step process which parcellates each subject’s cortex into 180 anatomically and functionally segregated regions per hemisphere. This first step involves the preprocessing and alignment of T1w and T2w scans, leveraging FreeSurfer to generate surface meshes. The pipeline then implements an algorithm called multimodal surface matching (MSM), in which a pre-trained classifer is applied to T1w, T2w, and fMRI scans to extract and fit the cortical surface to the 360 region HCP parcellation scheme (Glasser et al., 2016). In the case of HCP preprocessed datasets, the parcellation algorithm had already been run and the parcellated regions were extracted using connectome workbench.

However, not all datasets had undergone the HCP preprocessing pipeline, which is computationally expensive to run for large sample sizes and requires multimodal images that were not available for all datasets. For these datasets, we instead aligned images to the HCP atlas using T1w data alone. T1w scans were first preprocessed using FreeSurfer (version 7.4.1, (Fischl, 2012)) *recon-all* to obtain the cortical meshes. The Glasser atlas in fsaverage space was then applied to each subject’s surface mesh. The resulting atlas was then used to parcellate the freesurfer meshes. Absolute surface area (in *mm*^2^) for a parcel was calculated as the area sum over all vertices in the parcel. All surface mapping and parcellation steps were carried out in MATLAB version 9.7.0.1339299 R2019B.

Within each dataset, scans were sometimes acquired from different sites which can impact absolute structural measurements (Hedges et al., 2022). Therefore, ComBat harmonization was applied to the surface area values of dataset using the neuroCombat R package (Fortin, 2025). Scan site, which also encompassed preprocessing in those cases with participant differences, was used as the batch variable, while sex, age, and group of interest were included as covariates.

### 2.5 Structural covariance network construction

Surface area values for the 360 HCP parcels werefirst reduced to 180 values by taking the mean across hemispheres. SCNs are often defined using inter-regional Pearson correlations (Alexander-Bloch, Giedd and Bullmore, 2013); however, the weights of specific edges can be confounded by indirect relationships through other vertices (Masuda et al., 2018). Therefore, partial correlations were used instead, which regress out the effects of other vertices on specific pairwise relationships and have also been shown to better reflect structural connectivity (Joshi et al., 2010). This constructs a network in which each edge weight represents the unique shared variance in surface area between two regions.

Solving the partial correlation matrix via covariance matrix inversion was not possible due to rank deficiency (102 < 180). Instead of regularization methods, we opted for dimensionality reduction via hierarchical clustering (HC). To capture parcel dependencies, the HC algorithm, using Ward’s criterion (Ward, 1963), was run on the partial correlation matrix estimated from a sample of *N* = 650 healthy controls used in a prior study (Ward et al., 2024). To identify a set of robust clusters, numbers of clusters ranging from 9 to 100 were assessed for quality and consistency. Separation and compactness of clusters was measured using mean silhouette score and the Dunn index. Consistency of the clustering was measured using the adjusted Rand index, which was assessed for separate clusterings of data from the left and right hemispheres, as was as for clusterings from 20 random subsamples of *N* = 325 from the full *N* = 650 control group. These metrics were normalised and summed into a single score. To ensure the selected clustering was not due to biased random sampling, the process was repeated 10 times using 10 random seeds. This process suggested 62 clusters 8/10 times, and 57 clusters 2/10 times, so 62 clusters was selected as optimal. Clusters contained a range of 2-5 parcels, with a mean of 2.90. Intruigingly, 100% of clusters were comprised entirely of parcels that share borders. This finding suggests that neighbouring regions of the cortex share similar profiles of surface area relationships with the rest of the cortex. This is not to suggest that proximal regions necessarily have similar morphology, but that they expand or contract in a coordinated fashion with respect to the rest of the cortical surface.

Surface area values for each group were summed across parcels belonging to the same cluster. The partial correlation matrix 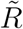 of the cluster SA values was estimated for each group via inversion of the covariance matrix Σ. To ensure partial correlations adequately captured the dependency structure of the clusters, cluster surface areas were statistically checked for normality using the Anderson-Darling test, as well as for differences in variance between compared groups using the F-test. Within each sample, the proportion of normally-distributed clusters was generally high, ranging from 60% *−* 90%, while the variance differences were zero or very low in most cases (Supplementary Material Table S1), suggesting that partial correlations were able to capture the majority of the dependence structure between the clusters. Graphs *G*(*V, E*) representing SCNs were created from the partial correlation matrices using the graph-tool python package (version 2.79, Peixoto (2017)), where the clusters defined vertices *v*_*i*_ (with *i* = 1, …, *n*, and with *n* = 62) and the absolute partial correlation magnitudes 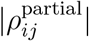 defined edge weights *w*_*ij*_ from vertex *v*_*i*_ to vertex *v*_*j*_. Weight thresholds were not applied, meaning SCNs were fully connected in the first instance. However, two subnetworks containing only positive or negative correlations were also analysed for each group.

### 2.6 Network metric calculations

Weighted network metrics including the global clustering coefficient, characteristic path length, global efficiency, and betweenness centrality were calculated using graph-tool. For path length, efficiency and betweenness centrality, edge weights were inverted prior to the calculation, as edge weights reflect association rather than cost. Global clustering *C*, which reflects the degree of organisation of network weights into triangles, is calculated according to Zhang and Horvath (2005) as:

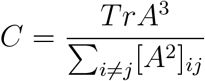

Where *A* is the connectivity matrix, in this case equal to the absolute partial correlation matrix 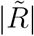. Characteristic path length *L*, representing the average shortest distance between vertices, was calculated as:

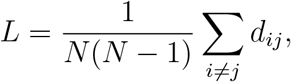

where each direct connection *A*_*ij*_ is treated as having length 1*/A*_*ij*_ and *d*_*ij*_ is the shortest-path distance between vertices *v*_*i*_ and *v*_*j*_ computed from these lengths. Similarly, global efficiency *E*_glob_, which reflects the ease of communication across the network, was calculated as:

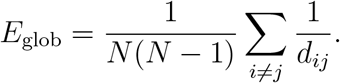

Betweenness centrality *C*_*B*_, which represents how many of the shortest paths travel via a given vertex *v*_*i*_, was calculated as:

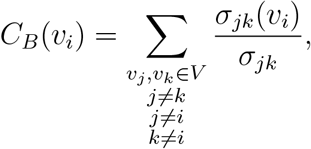

where *σ*_*jk*_ is the number of shortest paths from the source vertex *v*_*j*_ to target vertex *v*_*k*_ and *σ*_*jk*_(*v*_*i*_) is the number of shortest paths from *v*_*j*_ to *v*_*k*_ that pass through vertex *v*_*i*_. The values were normalised by the numbers of pairs of vertices not including *v*_*i*_.

Edge weight diversity was also investigated as a means of indexing network complexity. Jensen–Shannon (JS) divergence was used to capture heterogeneity in the patterns of edge weights between vertices, as this compares two probability distributions for their dissimilarity (Bai and Hancock, 2012). Distributions of edge weights *w* for each vertex *v*_*i*_ to all vertices *{v*_1_, …, *v*_*n*_*}* were first normalised to create a probability vector *p*_*i*_. For each specific target vertex *v*_*j*_:

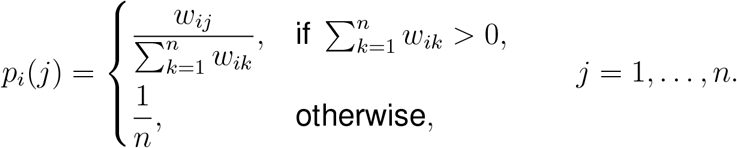

For any two vertices *v*_*i*_ and *v*_*j*_, let

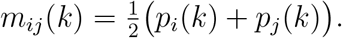

The prerequisite to JS divergence is the Kullback–Leibler (KL) divergence, which from from *p*_*i*_ to *m*_*ij*_ is calculated as:

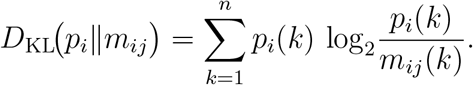

The JS divergence (JSD) between the edge weight distributions of vertices *i* and *j* is the symmetric version of the KL divergence and was calculated as:

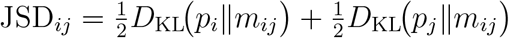

which satisfies JSD_*ij*_ = JSD_*ji*_ and JSD_*ii*_ = 0.

To establish the average divergence of one vertex *v*_*i*_, we took the mean of JS divergences of vertex *v*_*i*_ to all vertices:

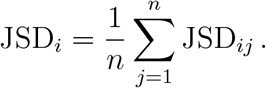

Finally, to establish a single JS divergence value for a network (JSD_global_), we took the mean JS divergence between all pairs of vertices:

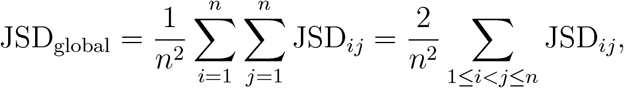

where the second equality uses the symmetry JSD_*ij*_ = JSD_*ji*_ and JSD_*ii*_ = 0.

In summary, JSD_global_ quantifies, in bits, the mean JS divergence between all pairs of edge weight distributions in the weighted network.

### 2.7 Cortical geodesic and small-world propensity calculations

Anatomically relevant distances between parcel clusters were estimated by calculating geodesics on the spherical cortical surface. Cluster centres were calculated as the 3-dimensional centroids of fsaverage vertices belonging to each cluster. This can be problematic in some cases when non-uniform region shapes lead to the centre of mass being outside the region itself. Therefore cluster centres were checked visually, and in all cases the centroids were located inside the clusters as the majority of clusters were roughly circular. The geodesic between centre points **P** of clusters *i* and *j* was calculated as:

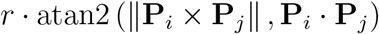

Where *r* is the sphere radius. The calculation was repeated for all pairwise combinations of clusters to obtain cortical distances associated with every cluster correlation.

Small-world propensity (SWP) was calculated as in Muldoon et al. (2016), using cortical distances between clusters as derived above. Empirical networks were reconfigured into two null models: a lattice network where the largest edge weights are at the shortest distances, and an ensemble of 10 random networks where edge weights were shuffled randomly. SWP *ϕ* was then calculated as a function of the observed and null global clustering coefficients and characteristic path lengths:

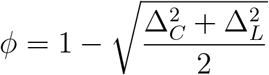

where

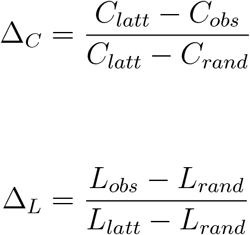

Delta *δ*, which represents the extent to which the divergence from perfect small-worldness (*ϕ* = 1) is due to a too lattice-like or too random-like topology, was calculated also as:

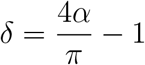

where

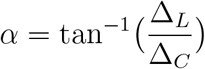

A delta close to −1 therefore relfects an observed network with a clustering coefficient maximally different to its lattice null model, while a delta close to +1 reflects an observed network with a path length maximally different to its random null model.2

### 2.8 Statistical analysis

Statistical analysis was carried in R (version 4.5.0, R Core Team (2025)) and python (version 3.1.0). Normality tested was performed using the Shapiro-Wilk test or the Anderson-Darling test. Group comparisons of distribution means and variances for normal distributions (e.g., partial correlations) were implemented with Welch’s t-test and F-tests repsectively. FDR correction was used in the case of sets of multiple tests.

Group comparisons of descriptive statistics for non-normal distributions (e.g., betweenness centrality) were implemented using permutation tests with 10,000 iterations. One-sided P-values were then calculated as the proportion of times the null group differences were more extreme than the observed group difference in the chosen test direction. Group comparisons of global network properties, for which only one value per network is obtained, were also implemented using permutation tests. Here, group labels were swapped at the level of the SA data and network metrics re-calculated within each iteration for a total of 10,000 iterations. Effect sizes for differences in global metrics were calculated as the observed group difference normalised by the standard deviation of the null difference distribution. Standard error for the global metrics was estimated using jackknife (leave-one-out) resampling.

The non-linear relationship between cortical geodesic distance and network weights was modelled using Generalized Additive Models (GAMs). The mgcv R package (Wood, 2017) was used to fit Gaussian GAMs with weight as the response variable, per-group distance smooth terms and a parametric group term. This process fits a smooth weight-distance curve separately for each group, and also estimates the overall difference in weight due to group. Smooth terms were always fit with a basis dimension *k* of 5, chosen during preliminary testing to provide adequate flexibility without overfitting. Otherwise, parameters were left as default. Statistical significance of the parametric and smooth terms was estimated via t-tests and approximate F-tests respectively. To test for differences in the group-specific smooths, the full model was compared with a reduced model lacking group-separated smooths using a *χ*^2^ analysis of deviance. To examine group differences along specific distances, model predictions at 100 equidistant points spanning the observed cortical distances were compared using pointwise Z-tests, applying FDR correction for multiple comparisons.

## 3 Results

### 3.1 Distributions of structural correlations

Partial correlations between cortical regions were normally distributed around zero in all cases (*P* >.05, *n* = 1891, Shapiro-Wilk test). To investigate potential differences in correlation structure between affected and control groups, we employed t-tests and F-tests. The t-tests were used to check for group differences in correlation sign, as a mean shift away from zero would indicate more positive or negative correlations on average. We found no significant differences in mean correlation, suggesting no biases towards positive or negative correlations in any condition. We then compared overall correlation magnitudes using F-tests for variances, as a change in variance would indicate correlations closer to or further from zero. These tests revealed significant increases in variance in the synaesthesia, ABIDE-I and -II datasets, as well as a significant decrease in the HCP-EP dataset (Table 2). This indicates that the synaesthesia and autism networks possessed greater magnitude correlations, while the early psychosis network had generally weaker correlations. Visual examination of the distributions suggested these effects applied for both positive and negative directions (Figure 2).

**Table 2:** Tests for group differences in partial correlation distributions. *n* = 1891 for all tests.

| Sample | t-test<br>$t_{(3740.8)}$ | t-test<br>P-value | F-test<br>$F_{(1890,1890)}$ | F-test<br>P-value |
| --- | --- | --- | --- | --- |
| Syn100 | 0.06 | .9489 | 0.79 | < .0001 |
| ABIDE-I | -0.03 | .9768 | 0.84 | .0001 |
| ABIDE-II | 0.03 | .9745 | 0.84 | .0001 |
| HCP-EP | 0.03 | .9778 | 1.10 | .0330 |
| BANDA | 0.03 | .9780 | 0.97 | .4517 |
| DCAM | 0.01 | .9910 | 0.96 | .4007 |
| PDC | 0.01 | .9894 | 0.98 | .6780 |

**Figure 1:**
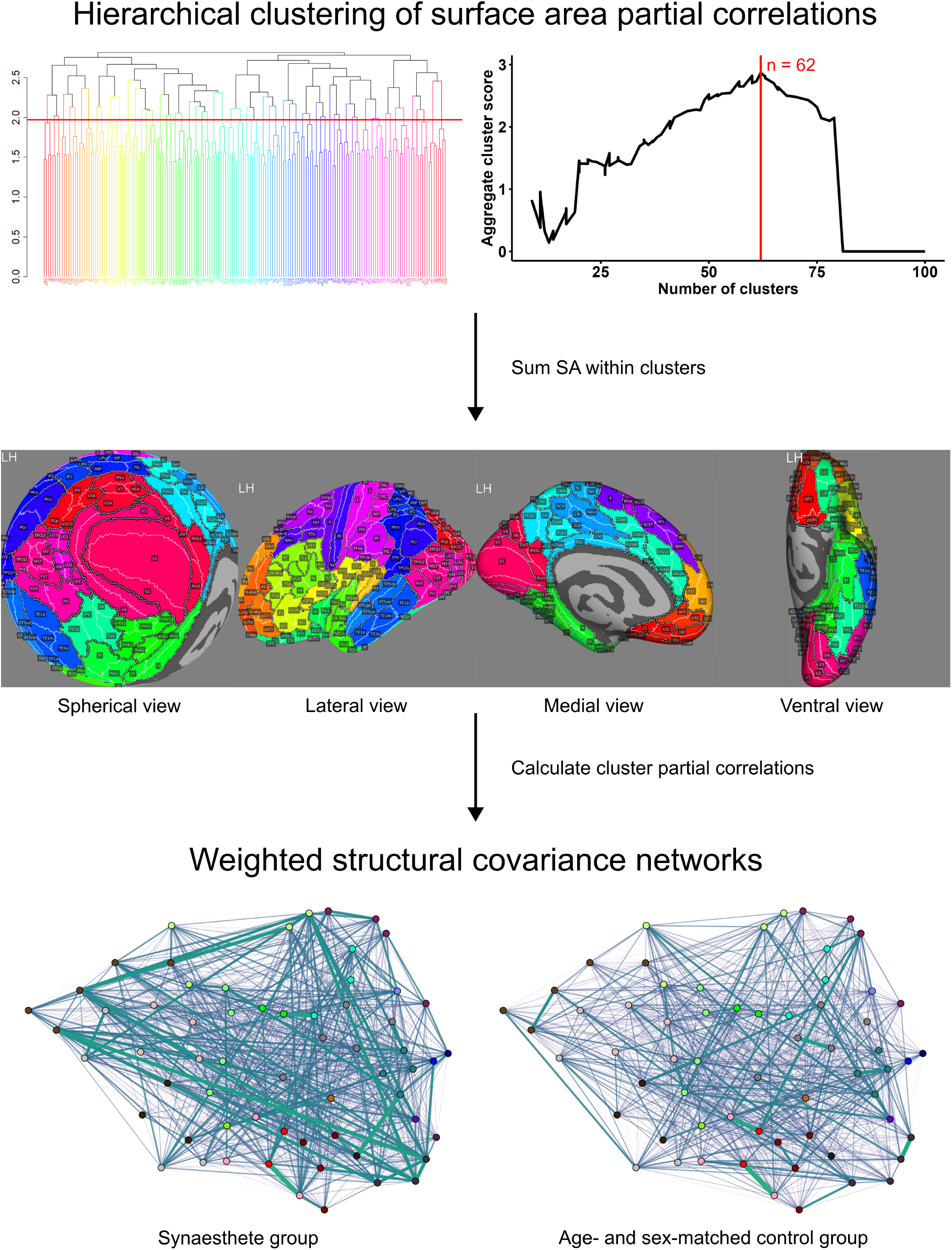
Process of constructing per-sample structural covariance networks (SCNs). Partial correlations between HCP parcel surface areas were calculated for a group of healthy controls (*N* = 650), which underwent hierarchichal clustering to generate 62 clusters of neighbouring parcels. Partial correlations between clusters were used generate SCNs for each control and affected group. In the SCN diagrams, line thickness and opacity reflects the magnitude of the partial correlation between two clusters.

**Figure 2:**
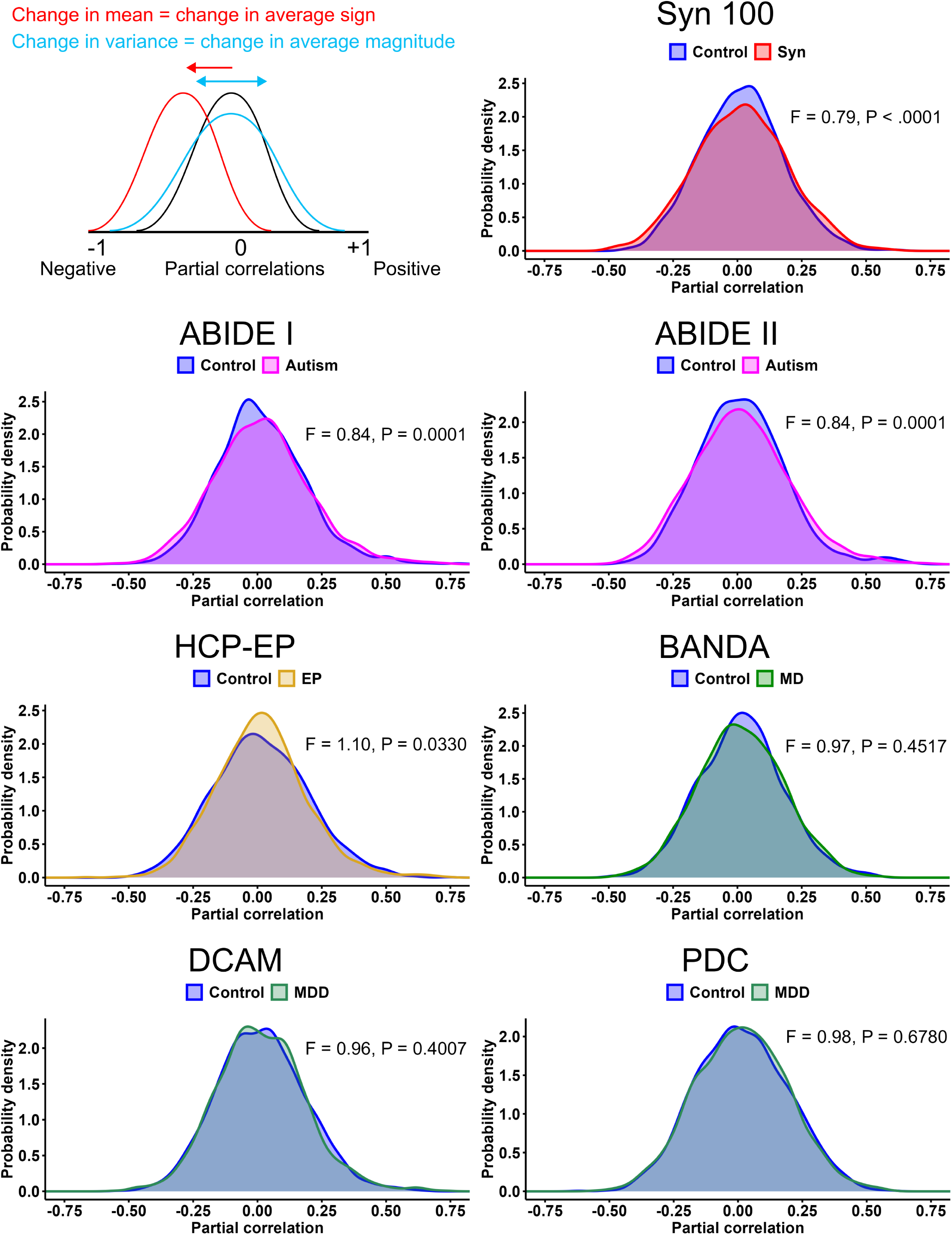
Distributions of partial correlations for control and affected groups in each dataset. Changes in mean would reflect changes in average sign, while changes in variance would reflect changes in average correlation magnitude. F-values pertain to F-tests for variance, *n* = 1891 observations.

### 3.2 SCN network properties

For the global network metrics, we hypothesised that synaesthesia and autism networks would display increased clustering and efficiency and reduced path length, in accordance with previous research using cortical thickness (Hänggi et al., 2011). In the synaesthesia network, a statistically significant decrease in path length was observed, while efficiency and clustering coefficient showed numerical but non-significant increases (*P* =.103). In ABIDE-I and ABIDE-II, the same pattern was observed, with some non-significant changes with the strongest effect for decreased path length in ABIDE-II. Following the diametric model (Crespi and Badcock, 2008), we hypothesised that HCP-EP would show opposing effects to autism and synaesthesia. However, while there were numerical decreases in efficiency and clustering, and an increase in path length, these were not statistically significant. In BANDA, DCAM and PDC, the same pattern as in synaesthesia and autism was observed, but there was no statistical significance (Figure 3, Table 3). The pattern of effect sizes follows the pattern of statistical significance, with relatively large effect sizes for the synaesthesia and ABIDE samples and smaller effects in the other samples. To check if these results were specific to the clustering scheme, the analysis for the Syn100 dataset was re-run across six adjustments to the number of clusters. The same patterns were observed for both the partial correlation distributions and network metrics (Supplementary Material Table S2), demonstrating these results were robust to region selection. Similar results were also observed in SCNs formed from positive or negative correlations only, with path length showing significant decreases in synaesthesia (*P* =.033) but non-significant decreases in autism (*P* =.059, Supplementary Material Tables S3 and S4). Overall, these results suggest that network topology in surface area SCNs may be best distinguished by characteristic path length. They also suggest that synaesthesia and autism possess similar network changes compared to controls.

**Table 3:** Group differences in global network metrics of SCNs. P-values were calculated via permutation tests with *n* = 10000 permutations. Effect size for each measure is analogous to Cohen’s *d*, calculated using the observed group difference in metrics normalised by the null (shuffled) distribution standard deviation

| Sample | Clustering | Clustering effect size | Clustering P-value | Global efficiency | Global efficiency effect size | Global efficiency P-value | Path length | Path length effect size | Path length P-value |
| --- | --- | --- | --- | --- | --- | --- | --- | --- | --- |
| Syn100 | Control: 0.136<br>Aff: 0.158 | 1.49 | .0645 | Control: 0.176<br>Aff: 0.196 | 1.64 | .0513 | Control: 3.119<br>Aff: 2.810 | -1.67 | **.0494** |
| ABIDE-I | Control: 0.139<br>Aff: 0.157 | 1.08 | .1262 | Control: 0.188<br>Aff: 0.204 | 1.24 | .1039 | Control: 2.932<br>Aff: 2.725 | -1.26 | .1033 |
| ABIDE-II | Control: 0.128<br>Aff: 0.137 | 0.99 | .1528 | Control: 0.172<br>Aff: 0.181 | 1.14 | .1264 | Control: 3.253<br>Aff: 3.083 | -1.2 | .1159 |
| HCP-EP | Control: 0.159<br>Aff: 0.147 | -0.75 | .2138 | Control: 0.200<br>Aff: 0.191 | -0.76 | .2188 | Control: 2.759<br>Aff: 2.911 | 0.94 | .1737 |
| BANDA | Control: 0.140<br>Aff: 0.142 | 0.1 | .4565 | Control: 0.179<br>Aff: 0.183 | 0.34 | .3654 | Control: 3.088<br>Aff: 3.005 | -0.46 | .3204 |
| DCAM | Control: 0.147<br>Aff: 0.147 | 0.01 | .5099 | Control: 0.187<br>Aff: 0.191 | 0.31 | .6273 | Control: 2.948<br>Aff: 2.904 | -0.25 | .6003 |
| PDC | Control: 0.156<br>Aff: 0.157 | 0.03 | .5091 | Control: 0.192<br>Aff: 0.195 | 0.25 | .5974 | Control: 2.875<br>Aff: 2.833 | -0.23 | .5902 |

**Figure 3:**
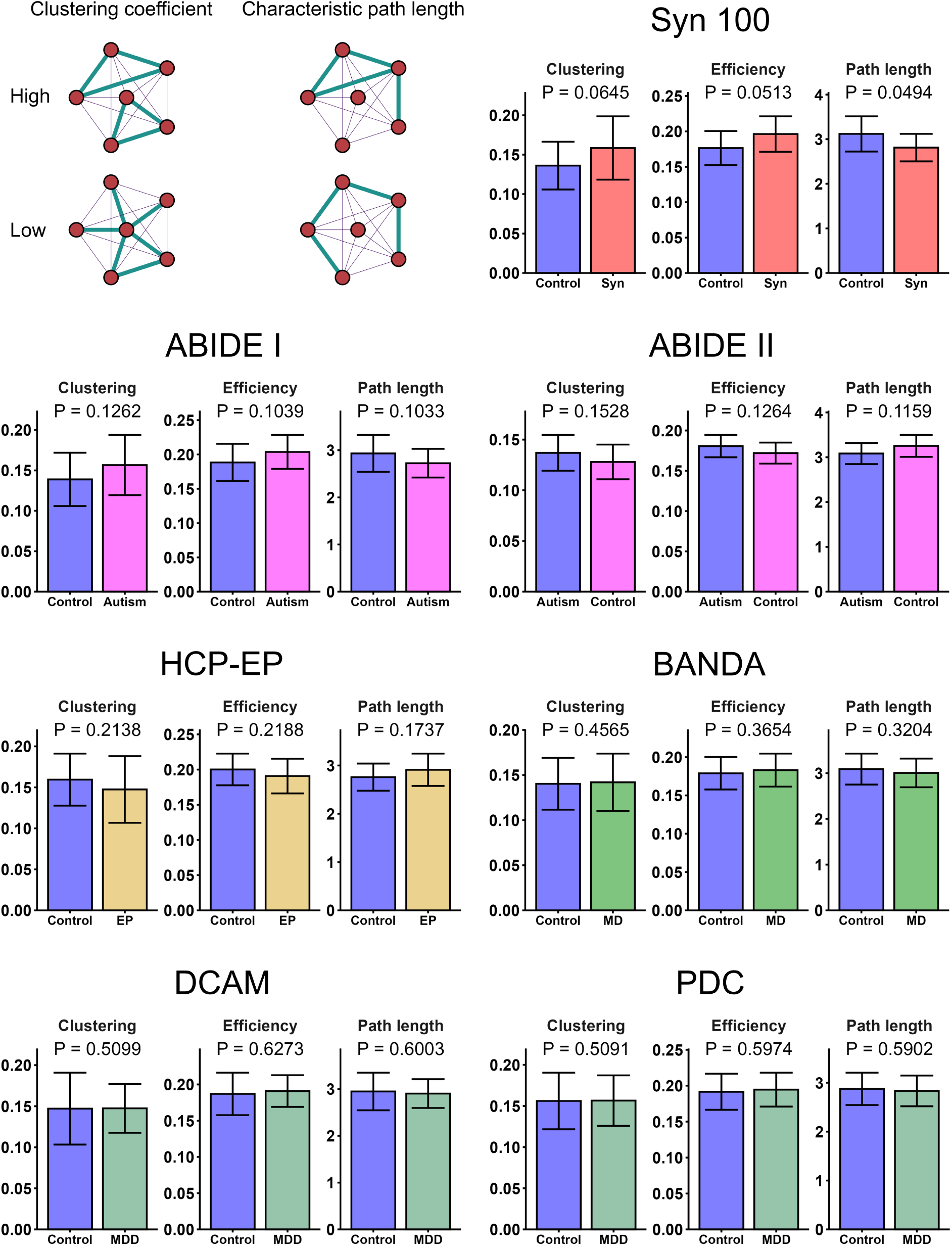
Global network metrics of SCNs. The diagram in the top left shows toy networks with high and low clustering and path length, whereby the thicker lines represent larger edge weights. P-values were calculated via permutation tests with 10, 000 permutations, while standard error was estimated through jackknife resampling.

The distribution of centrality values across vertices can also provide insight into the hierarchical organisation of the network. All samples displayed a centrality distribution characterised by many vertices with low centrality and decreasing numbers of vertices with higher centrality. In line with research on functional connectivity (Hong et al., 2019; Ward et al., 2024), we hypothesised that synaesthesia and autism networks would show compressed hierarchies compared to their controls, as indicated by flatter distributions of centrality values. However, except for a small increase in variance in synaesthesia, comparison of distribution shape revealed no group differences in any datasets (Figure 4). The small effects observed here suggest that none of these conditions are associated with a departure of the structural network from the typical hierarchical organisation characterised by many low-importance regions and fewer high-importance regions. Similar results were observed in positive- and negative-only networks (Supplementary Material Tables S5 and S6). Furthermore, across samples, mean group differences in correlation strength showed no clear differences between known functional connectivity networks (Yeo et al., 2011), suggesting global structural changes were also not predominantly localised within specific functional regions (Supplementary Material Table S7).

**Figure 4:**
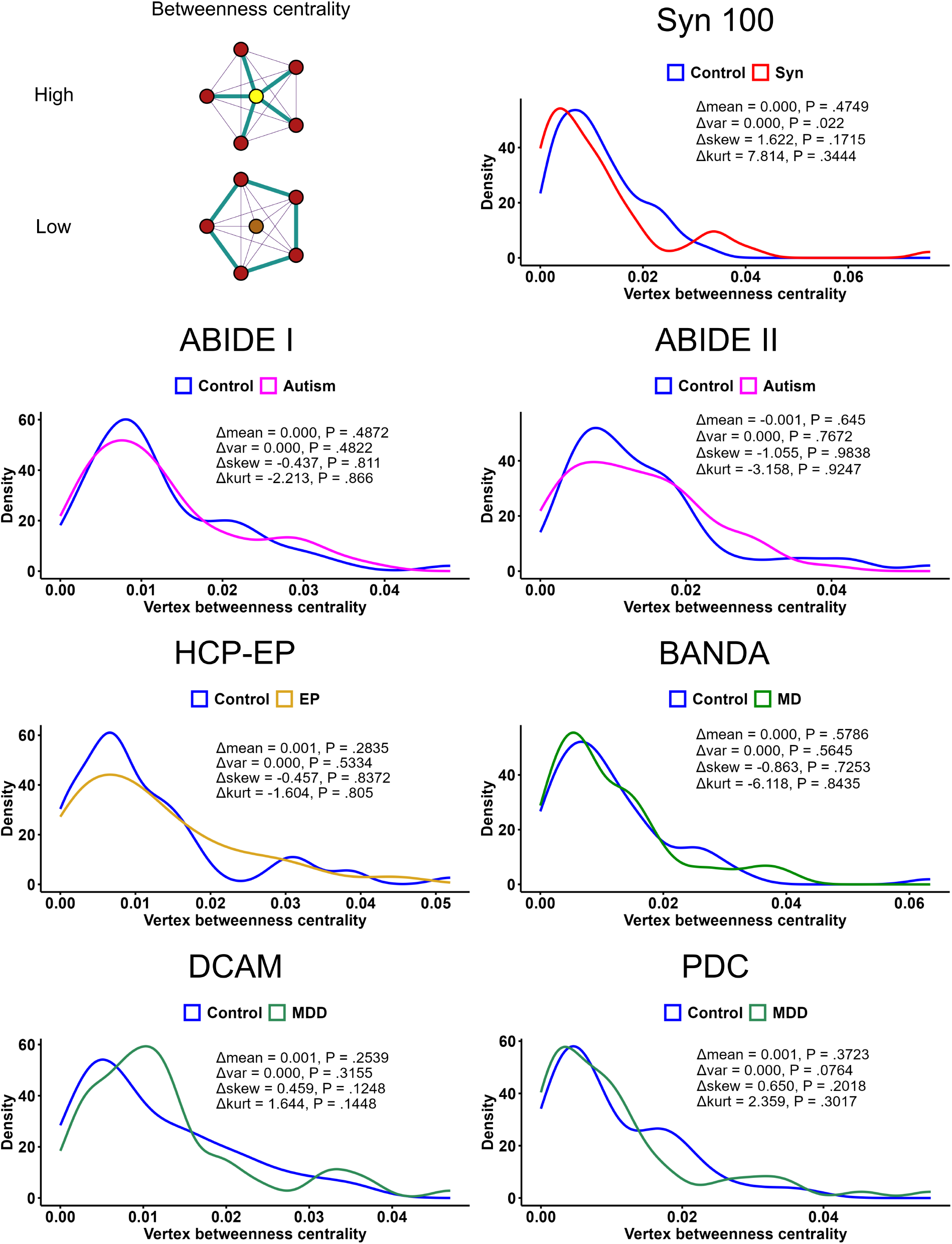
Distributions of betweenness centrality values. The diagram shows toy networks where the central vertex has high and low betweenness centrality. For each dataset, betweenness values were compared between affected and control groups in terms of their Pearson correlation, means, variance, skewness and kurtosis. P-values were estimated via permutation tests for 10, 000 permutations, *n* = 62 observations.

Network complexity was assessed using Jensen–Shannon divergence between the edge weight distributions of all vertices. Significant decreases in mean JS divergence were observed in synaesthesia, ABIDE-I and -II, and BANDA samples, as well as a significant increase in the HCP-EP sample, the latter showing the largest effect size (Table 4). This suggests that synaesthesia, autism and adolescent anxiety SCNs have decreased complexity, while the early psychosis SCN bears substantially increased complexity. These differences could further reflect the globally increased cortical interactivity in synaesthesia and autism, leading to more homogeneous associations between regions, while the reduced interactivity in early psychosis reflects regions with more specialised associations.

**Table 4:**
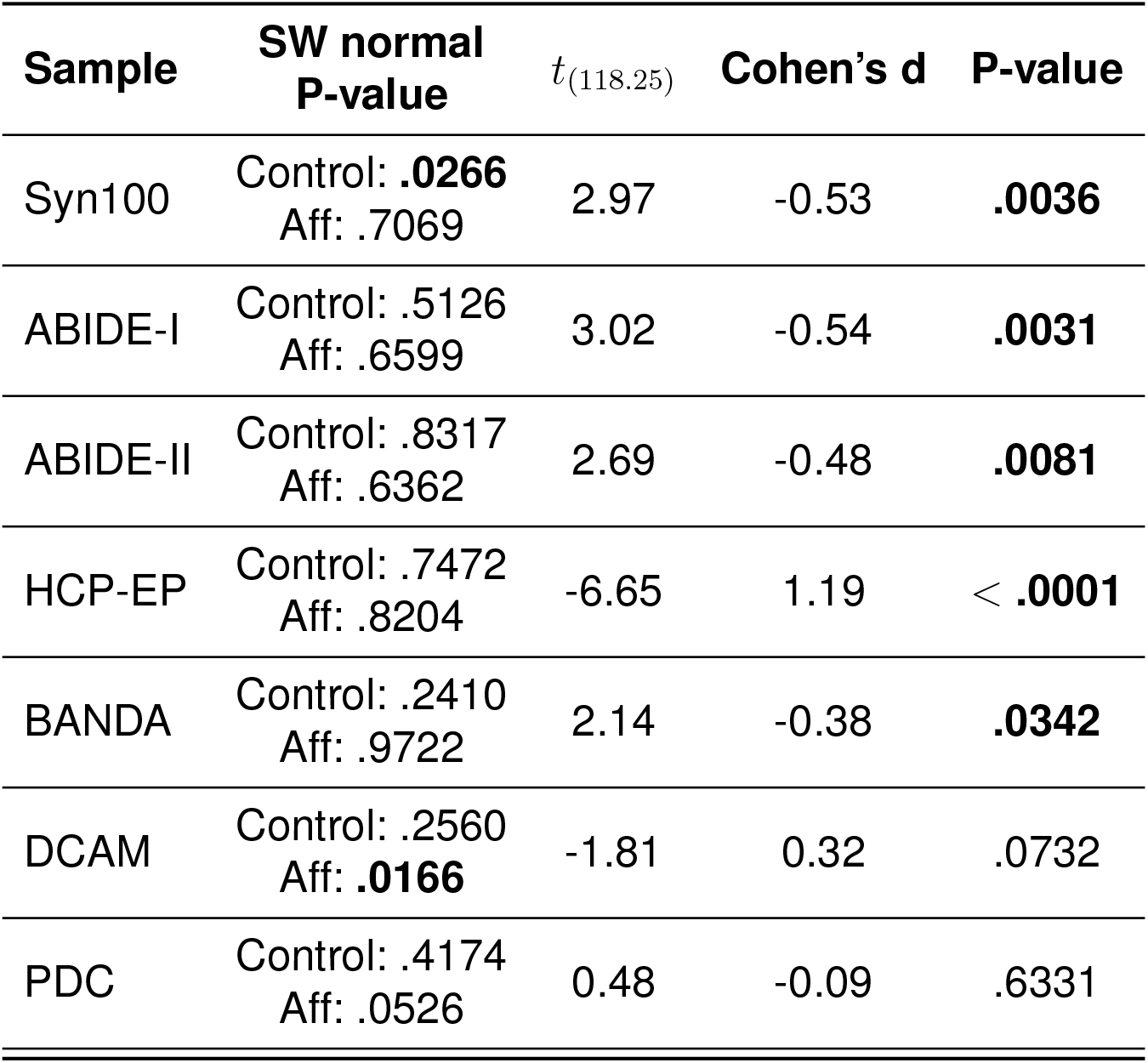
Group differences in mean Jensen-Shannon divergence of edge weight distributions. *n* = 62 observations. SW = Shapiro-Wilk.

### 3.3 Spatial organisation of network weights

To assess if changes in network weights followed a spatial pattern, we modelled the relationship between weight and cortical distance using generalized additive models (GAMs). We hypothesised that autism and synaesthesia would have stronger weights at longer distances over the cortex, in line with structural and functional connectivity findings in autism (Müller et al., 2011; Weber et al., 2024). All samples displayed a highly significant smooth term for distance (Table 5), indicating that distance has a consistently non-linear relationship with network weights. This relationship was characterized by strong weights at short distances that quickly flattened to consistent yet lower weight at longer distances (Figure 5), suggesting that like other brain networks, SCNs possess the strongest weights at short distances. In no case was a significant interaction between group and the smooth distance term observed (Table 5), suggesting that the conditions analysed do not deviate from this regular pattern. Significant effects of group were observed for the synaesthesia, ABIDE-I and -II and HCP-EP datasets, but not for the DCAM, PDC or BANDA datasets (Table 5). This provides further support for the hypothesis that synaesthesia, autism and early psychosis show differences in global correlation strength, but that anxiety and depression groups have overall similar correlation magnitudes to healthy controls.

**Table 5:** Generalised additive models (GAM) predicting network weight as a function of a parametric Group term and separate smooth Distance terms for each group. Estimated and reference degrees of freedom for F values can be found in Supplementary Material Table S10. *n* = 1891 observations per group.

| Sample | Group coefficient | Group coefficient SE | Group coefficient $t_{(3772.363)}$ | Group coefficient P-value | Smooth term F | Smooth term P-value | Group:smooth deviance | Group:smooth P-value |
| --- | --- | --- | --- | --- | --- | --- | --- | --- |
| Syn100 | 0.016 | 0.003 | 4.918 | < .0001 | Control: 24.21<br>Aff: 25.34 | Control: < .0001<br>Aff: < .0001 | 0.032 | .5289 |
| ABIDE-I | 0.013 | 0.003 | 3.836 | .0001 | Control: 75.12<br>Aff: 84.43 | Control: < .0001<br>Aff: < .0001 | 0.061 | .2295 |
| ABIDE-II | 0.014 | 0.003 | 4.037 | < .0001 | Control: 91.32<br>Aff: 70.56 | Control: < .0001<br>Aff: < .0001 | 0.045 | .3791 |
| HCP-EP | -0.011 | 0.003 | -3.312 | .0009 | Control: 57.10<br>Aff: 63.99 | Control: < .0001<br>Aff: < .0001 | 0.038 | .4857 |
| BANDA | 0.004 | 0.003 | 1.147 | .2515 | Control: 25.86<br>Aff: 18.70 | Control: < .0001<br>Aff: < .0001 | 0.038 | .4051 |
| DCAM | 0.000 | 0.003 | 0.144 | .8855 | Control: 41.17<br>Aff: 57.88 | Control: < .0001<br>Aff: < .0001 | 0.061 | .2032 |
| PDC | 0.001 | 0.003 | 0.219 | .8267 | Control: 19.76<br>Aff: 14.88 | Control: < .0001<br>Aff: < .0001 | 0.049 | .3242 |

**Figure 5:**
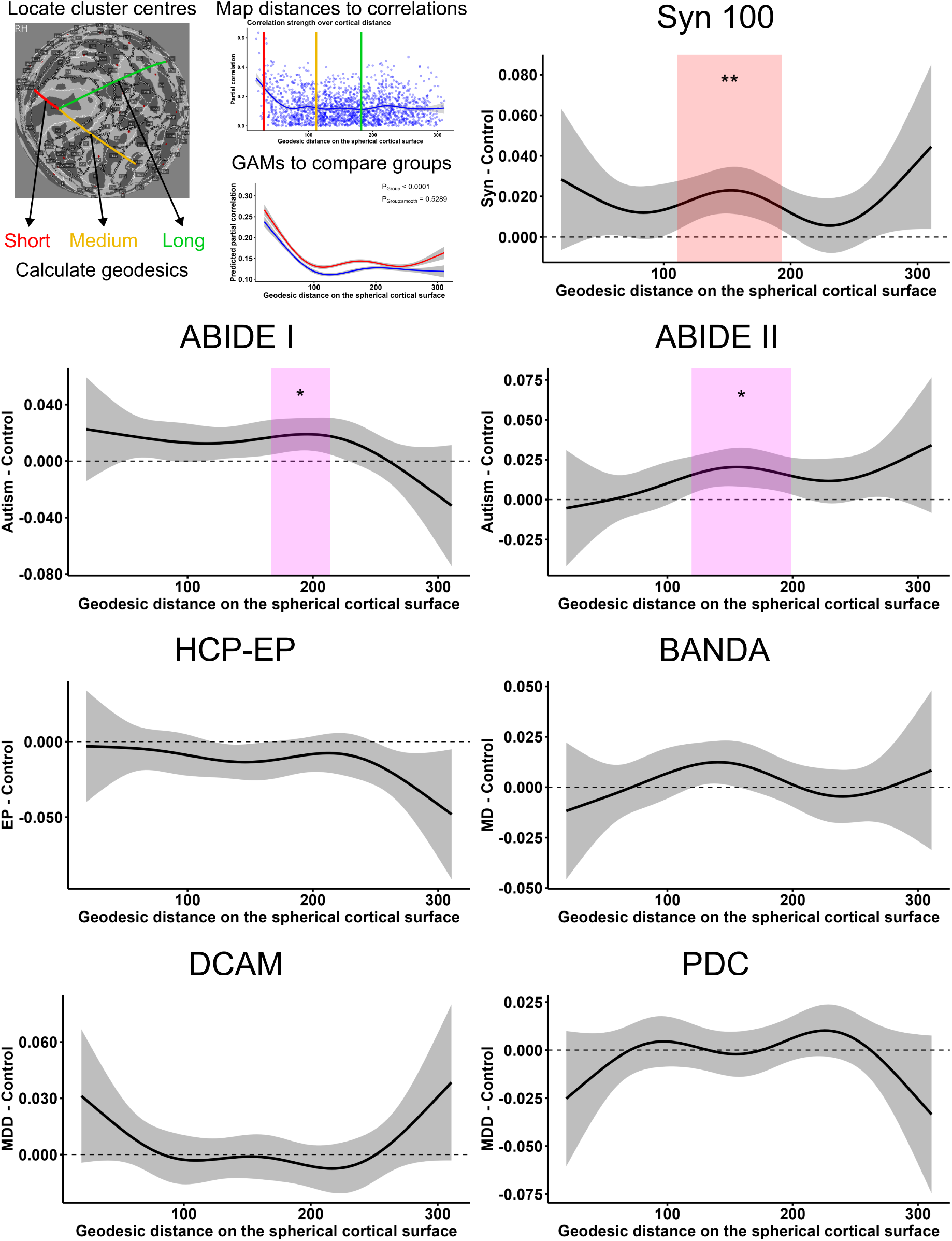
Group differences in network weights over the range of inter-cluster cortical distances. GAMs were used used to model the smooth relationship between cortical distance and network weight, according to group. The plots for each dataset show the group difference in predicted network weights across the range of cortical distances, with the shaded area indicating the 95% confidence interval. Distances with significant group differences are highlighted; ‘**’ = *P* <.01, ‘*’ = *P* <.05, with FDR correction for 100 tests. GAMs were fitted with *n* = 1891 observations per group.

When comparing differences in weight between affected and control groups across varying distances, significant differences were observed for the Syn100, ABIDE-I and ABIDE-II networks, which all had increased correlation strength at moderate distances (Figure 5). Therefore it appeared that in addition to globally increased correlation strength, synaesthesia and autism were further characterised by specific increases at moderately long cortical distances compared to controls. In contrast, early psychosis showed no significant differences at specific cortical distances, suggesting a more distributed decrease in correlation strength. Somewhat similar results were observed in SCNs with positive- or negative-only correlations. Interestingly however, only positive SCNs showed the non-linear relationship between distance and weight, while negative SCNs showed no clear relationship (Supplementary Material Tables S9 and S10). Furthermore, synaesthesia showed significant weight differences at moderate- and very-long distances in the positive network, but also differences at short-to-moderate distances in the negative network; in contrast, autism showed significant differences at moderate distances in the negative network only (Supplementary Material Figures S1 and S2).

Spatial information was also analysed by calculating small-world propensity (SWP), which compares the observed network to lattice and random reconfigurations. Because of the computational cost of the SWP calculation, we were unable to reliably estimate statistical significance via permutation tests which were computationally unfeasible. Therefore, we report the SWP and delta measurements but make no conclusions about the robustness of any group differences. Across datasets, we observed small-world propensity (SWP) values for controls in range of 0.4-0.6 (Table 6, Figure 6), which is relatively typical if not slightly low for brain networks (Muldoon et al., 2016). In the synaesthesia and autism networks, SWP values were reduced compared to controls with an absolute difference of 0.1. Conversely, SWP was slightly increased in the HCP-EP, BANDA, DCAM and PDC datasets, although this was very small for the latter sample. This suggests that synaesthesia and autism SCNs could be characterised by reduced small-worldness. Examination of delta values, which were all close to *−*1, indicated that all the SCNs are characterised by very random-like topology.

**Table 6:**
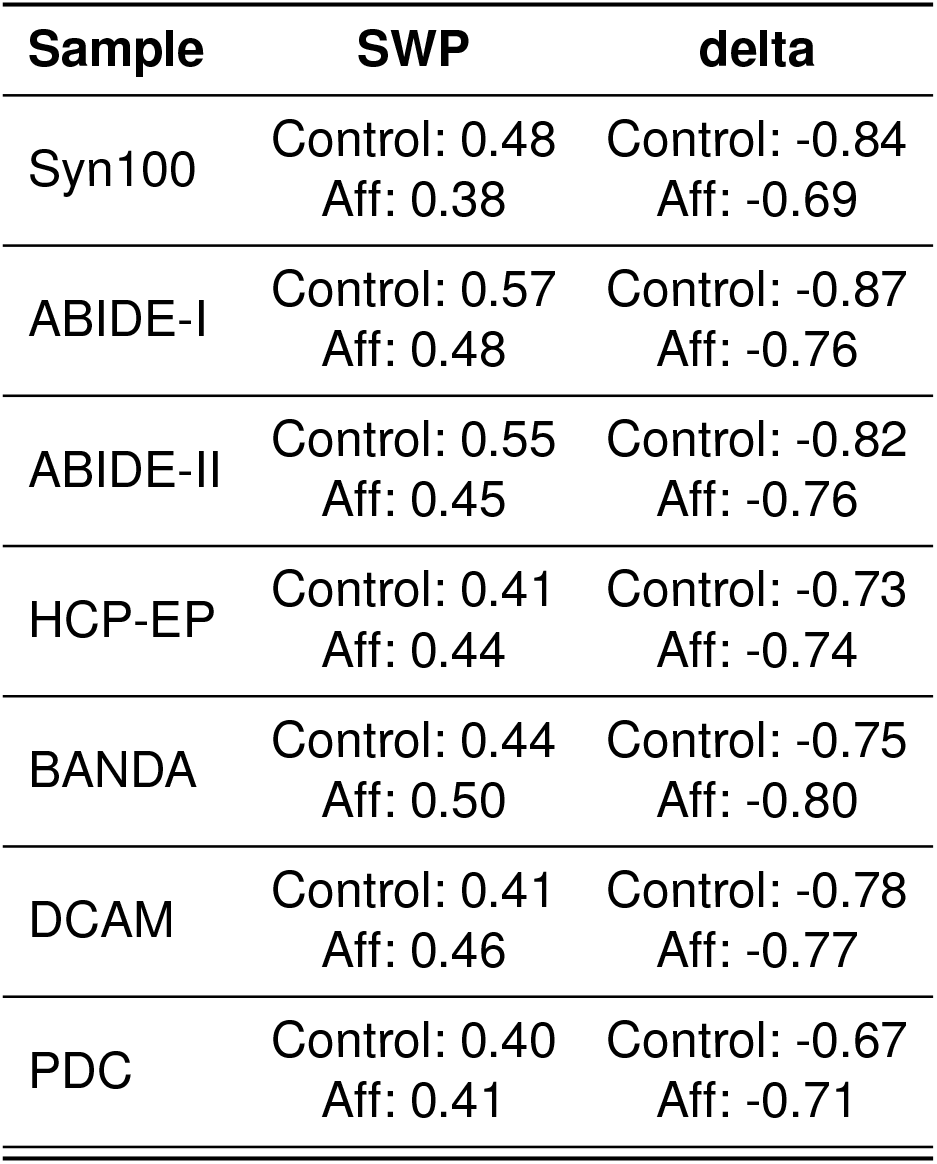
Small-world propensity (SWP) and delta values of SCNs.

| Sample | SWP | delta |
| --- | --- | --- |
| Syn100 | Control: 0.48<br>Aff: 0.38 | Control: -0.84<br>Aff: -0.69 |
| ABIDE-I | Control: 0.57<br>Aff: 0.48 | Control: -0.87<br>Aff: -0.76 |
| ABIDE-II | Control: 0.55<br>Aff: 0.45 | Control: -0.82<br>Aff: -0.76 |
| HCP-EP | Control: 0.41<br>Aff: 0.44 | Control: -0.73<br>Aff: -0.74 |
| BANDA | Control: 0.44<br>Aff: 0.50 | Control: -0.75<br>Aff: -0.80 |
| DCAM | Control: 0.41<br>Aff: 0.46 | Control: -0.78<br>Aff: -0.77 |
| PDC | Control: 0.40<br>Aff: 0.41 | Control: -0.67<br>Aff: -0.71 |

**Figure 6:**
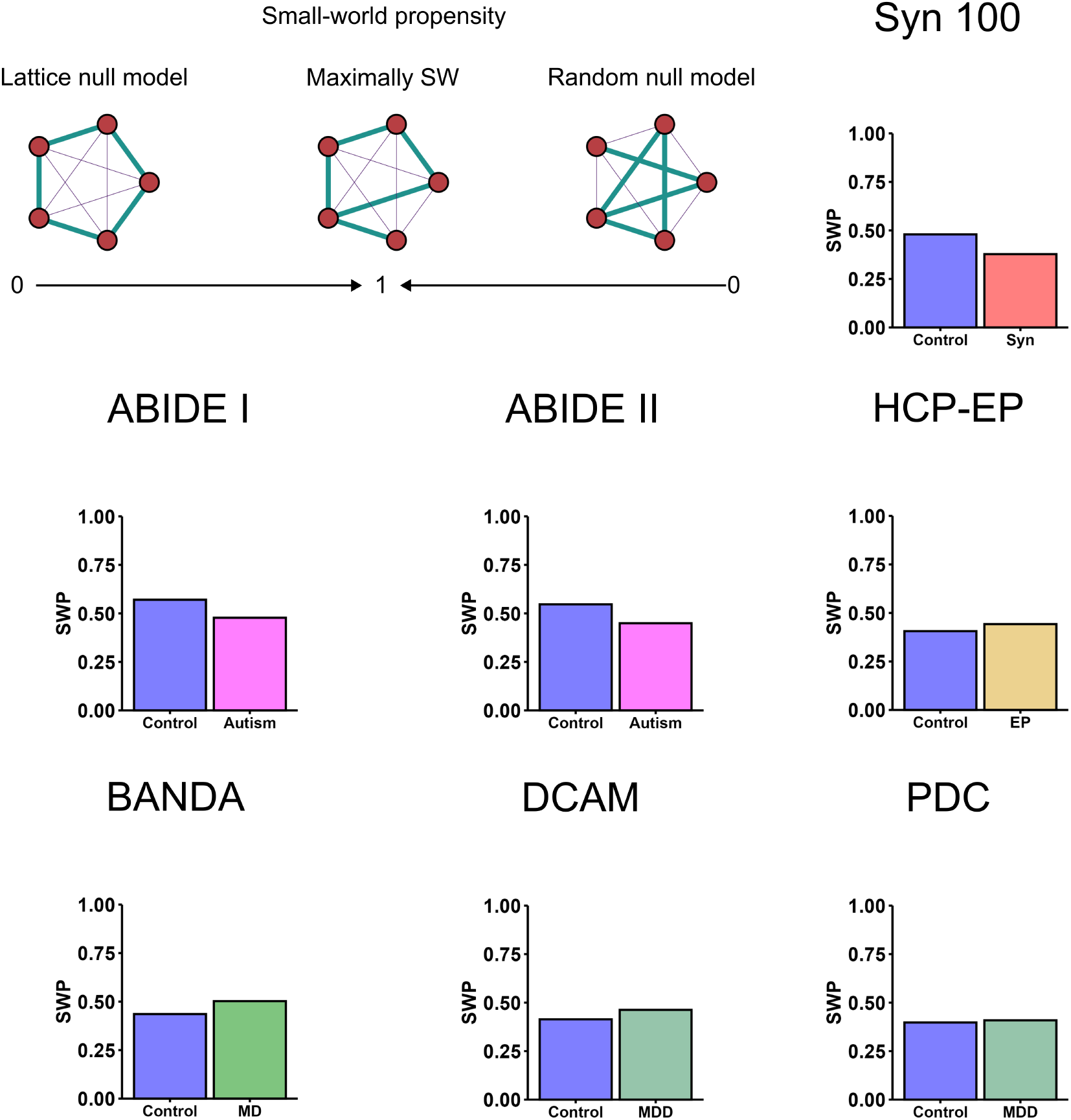
Small-world propensity (SWP) values. Small-worldness is largest when a network has a balance short path length, as in a random network, and high clustering coefficient, as in a lattice network. SWP was calculated by comparing the observed SCNs to null models where edge weights were randomised (random) and where the edge weights were ordered from largest to smallest over increasing cortical distance.

Interestingly, delta values were more positive in the synaesthesia and autism networks compared to their controls, suggesting these networks were closer to a lattice configuration (i.e. strong short range connectivity) than control networks despite having reduced SWP (Table 6,Figure 6). This apparent discrepancy is due to the fact that SWP was calculated for each SCN with reference to its own null models: therefore, synaesthesia and autism SCNs were further from both their own lattice and random null models, but slightly closer to their lattice null model compared to control networks. The decrease in SWP in synaesthesia and autism may be explained by the increases in correlation magnitudes at moderate-range distances in these networks, which confers neither local clustering nor shorter path lengths.

For positive and negative-only SCNs, findings diverged somewhat. The pattern of results was consistent in the negative SCNs, although SWP was generally lower. In contrast, all positive SCNs had the same SWP value of *~* 0.7 (Supplementary Material Table S11). This suggests that the small-world properties of SCNs is driven primarily by the positive correlations, and this is invariant across populations.

## 4 Discussion

The structural organisation of the brain reflects neurodevelopmental processes that underlie the coordinated maturation of distributed regions, which shapes both their specialisation and integration (Miterko et al., 2018; He et al., 2025). Structural covariance networks (SCNs) provide a window into this organisation by describing how variation in one region is linked to variation in others. This is assumed to reflect the influence of a cross-regional latent mechanism during brain development. Notably, the degree of structural covariance between two regions within a particular demographic is associated with resting-state function connectivity, white matter connectivity, and mutual genetic influences, offering insights into how regions of the cortex interact (Schmitt et al., 2008; Alexander-Bloch, Giedd and Bullmore, 2013; Alexander-Bloch, Raznahan, Bullmore and Giedd, 2013; Moura et al., 2017). However, through the assessment of global network topology, cortical SCNs also provide an opportunity to capture higher-order organisational properties like efficiency, hierarchy, small-worldness and entropy (He et al., 2009; Berlot et al., 2016; Kim et al., 2016), and these properties may be more relevant for high-level experiential traits (Mišić and Sporns, 2016; Tompson et al., 2018). By collapsing group-wide cortical organisation to single metrics, this approach can also help mitigate statistical confounds commonly present in brain-wide association studies (BWAS) that examine properties of many areas or connections (Marek et al., 2022; Burns et al., 2025). Despite this advantage, some SCN investigations of conditions like autism have failed to replicate across studies (Bethlehem et al., 2017; Cai et al., 2021; Wang et al., 2022), likely due to methodological discrepancies (Carmon et al., 2020). Thus there is a need to examine several populations under the same framework (Ward, 2021).

In this study, SCNs were leveraged to characterise the organisation of cortical surface area in several populations with atypical experiences, as compared to age- and sex-matched controls.

Hierarchical clustering of the whole-cortex HCP atlas revealed that neighbouring regions possessed similar expansion-contraction relationships with the rest of the cortex. Distributions of partial correlations beween the region clusters revealed significantly stronger associations in synaesthesia and autism, weaker correlations in early psychosis, and no change in anxiety or depression. Across SCNs using absolute, positive or negative correlations, characterisation of network topology revealed significantly lower path lengths in synaesthesia, as well as a non-significant lower path lengths in autism. Although changes in early psychosis were in the opposite direction, these were not statistically significant. Autism, synaesthesia, and adolescent anxiety networks were also characterised by significantly decreased network complexity, as measured by the divergence of edge weight distributions across vertices, while the opposite pattern was observed in early psychosis. To determine if changes in network connectivity were more prominent at specific distances, we also modelled network weights as a function of their cortical distance. Synaesthesia and autism were both characterised by increases in correlation strength at moderate distances, with diverging patterns for positive and negative correlations. Finally, the degree of small-worldness was also smaller in synaesthesia and autism networks, and slightly larger in early psychosis, anxiety and depression networks, although this was not statistically tested. The results for autism and depression were replicated across two datasets, suggesting the analysis was robust.

Our results for synaesthesia are relatively consistent with a previous study examining cortical thickness correlations in 24 grapheme-colour synaesthetes, which found decreased path length, increased global clustering, and decreased small-world properties (Hänggi et al., 2011). This is reassuring, since in addition to the different structural measure and smaller sample, this study also made disparate methodological choices including different cortical atlases, Pearson correlations, and threshold application (Hänggi et al., 2011). Therefore, the finding of increased structural network connectivity in synaesthesia appears robust. SCN studies for autism and schizophrenia or psychosis are more numerous, but also more difficult to assess; in contrast to our global approach, they have typically focussed on the covariance of *a priori* selected regions, used seed-based networks, applied strong thresholding, or employed other methodological constraints that make direct comparison unfeasible (Prasad et al., 2022). Yet, some studies have used a global approach. A study using cortical thickness correlations from autistic children saw decreased network modularity and stronger inter-module connectivity (Shi et al., 2013), which broadly aligns with our findings of decreased complexity and globally increased correlation strength. A more recent study of cortical thickness and surface area SCNs in autistic infants also found decreased path length and increased global efficiency compared to controls, yet also decreased global clustering (Wang et al., 2022). Similarly to our approach, Bethlehem and colleagues also examined the relationship between correlation strength and cortical distance in autism using the ABIDE-I dataset. While this study found increases in correlation of surface area at moderate distances, they reported increases correlations of cortical thickness at short distances and decreases at longer distances. However, methodological details could explain this divergence: in addition to using cortical thickness rather than surface area, they measured distance as Euclidean distance between centroids (distance through the brain) while this study used geodesics (distance around the brain), and they modelled the distance-weight relationship linearly whereas we used a non-linear approach. Although they conclude a decrease in overall covariance in autism, this was only reflected in high-degree vertices (i.e., hubs) (Bethlehem et al., 2017). For psychosis and schizophrenia, studies that have examined global properties of SCNs generally agree with the current pattern of results for early psychosis, collectively finding longer path length, and lower integration, efficiency, and correlation strength in networks based on cortical thickness and white matter connectivity (Zhang et al., 2012; Griffa et al., 2015; Kim et al., 2020). Therefore, our finding of globally decreased correlation magnitude in early psychosis appears unlikely to be spurious, and perhaps reflects an early manifestation of a global schizophrenia brain phenotype. Previous results for global SCN topology in major depression have been mixed, with at least two studies finding increases in randomness reflecting a lower clustering coefficient and increased integration (Wang et al., 2016; Ping et al., 2023), and one study showing reduced global efficiency (Chen et al., 2022). Differential and null findings in depressed cohorts could reflect that the structural correlates of depression in the brain are likely to be highly individual (Han et al., 2023).

A major benefit of the current study is being able to contrast findings across these different populations. We conclude that our observed group differences cannot be due to methodological variability unlike previous studies, where groups have been studied separately with different methodological choices. Aside from population heterogeneity, another explanation for the null results in anxiety and depression is that these conditions are driven more by environmental than neurodevelopmental factors. Autism and schizophrenia are both highly heritable with a heritability around 0.7 (Tick et al., 2016; Hilker et al., 2018), and there is evidence to suggest that synaesthesia is also highly driven by genetic factors (Bosley and Eagleman, 2015), although some evidence suggests that synaesthesia a manifestation of a broader inherited cognitive profile, i.e. a ‘synaesthestic disposition’ (Ward and Filiz, 2020). Although depression and anxiety are thought to have a genetic component with a heritability around 0.3 (Sullivan et al., 2000; Hettema et al., 2001), their manifestation is largely dependent on social circumstances, trauma and adversity. Thus while they are associated with structural changes such as cortical thinning (Kim et al., 2019; Schmaal et al., 2020), this may not translate to robust patterns of covariance differences. Indeed, the search for a consistent biological signature of anxiety or depression may be fundamentally flawed because these manifestations reflect individual reactions to external events rather than a genetically-driven neurodevelopmental process (Brückl et al., 2020). Thus, our results across populations are in agreement with the notion that SCNs reflect neurodevelopmental (and aging) patterns more strongly than experience-dependent plasticity or instantaneous functional connectivity (Geng et al., 2017; DuPre and Spreng, 2017).

The finding of similar global SCN changes in autism and synaesthesia aligns with the hypothesis that these conditions share a common or overlapping neurodevelopmental process (van Leeuwen et al., 2020; Nugent and Ward, 2022). In particular, globally increased connectivity, decreased complexity and lower small-worldness suggest a shift towards a more random structural organisation, a conclusion also reached by others researching autism (Ouyang et al., 2022; Wang et al., 2022). This also converges with functional studies, which indicate globally increased connectivity and contracted functional gradients, indicating increased communication between sensory and association cortices in autism (Müller et al., 2011; Hong et al., 2019; Ilioska et al., 2023; Lee et al., 2025). These converging neurobiological results may also align with theoretical predictive processing accounts of autism that suggest an increased weight on sensory information (Lawson et al., 2014; Van de Cruys et al., 2017; Friston, 2020), in that increased cortical associations could reflect a heightened tendency to pass sensory information up the cortical hierarchy. A more outstanding question is how synaesthesia and autism are differentiated, given their co-occurrence is heightened but not complete. Although we find broadly similar results for these conditions, they diverged on the role of correlation sign at different distances: synaesthesia showed greater strength positive correlations over moderate and long distances, as well as greater strength negative correlations at short-moderate distances, while autism only showed greater strength negative correlations at moderate distance. Since positive structural correlations have been shown to reflect white matter connectivity significantly more robustly than negative correlations (Gong et al., 2012), this difference could reflect varying aspects of increased connectivity across synaesthesia and autism, which are likely to co-occur under neurodevelopmental regimes that increase connectivity more generally. Our cross-condition analysis also showed that early psychosis has reduced structural correlations as well as increased network complexity. The latter finding in particular aligns with studies showing increased hiearchical organisation in schizophrenia (Yang et al., 2023; Acero-Pousa et al., 2025), but also contrasts with studies showing greater homogeneity of connectivity (Mastrandrea et al., 2021). This current contrast between autism and psychosis finding lends itself to the diametric model of autism and psychosis, which suggests that these two conditions reflect opposing trade-offs in cognitive profile, with autism lacking mentalising ability and psychosis involving hyper-mentalising (Crespi and Badcock, 2008; Rządeczka et al., 2023). Although this model has received recent support from neurobiological findings showing disparities in EEG responses (Tarasi et al., 2022, 2023), more research is needed to test if this is also observed in global network topology.

### 4.1 Limitations

The study presents several important limitations. Our sample size was limited to *N* = 102, as this was the size of the smallest sample (Syn100), and we wanted to facilitate equivalent analyses across the datasets. Although this sample size is notably larger than most neuroimaging studies of synaesthesia (Racey et al., 2023), it may have lead to low statistical power across all samples in this study. Given the highly consistent patterns in network properties across datasets involving similar populations (ABIDE-I and ABIDE-II, or BANDA, DCAM and PDC), it appears more likely that lack of statistical significant differences in network metrics, particularly for autism, was driven by low statistical power rather than a true null effect. Future studies of this kind may benefit from pooling data from several samples to achieve larger sample sizes.

Relatedly, the use of hierarchical clustering to deal with the sample size being less than the number of HCP parcels (rank deficiency) also presents specific issues. On one hand, hierarchical clustering is a commonly-used approach for constructing cortical atlases or identifying networks based on voxel-wise features (Moreno-Dominguez et al., 2014; Doucet et al., 2019). For partial correlation estimation, it is easier to interpret than commonly used regularization methods, such as graphical lasso (Friedman et al., 2008) or Ledoit-Wolf regularization (Brier et al., 2015), since it preserves empirical relationships, while the latter approaches can have unpredictable effects on the dependency structure. However, HC inherently decreases atlas resolution, which is known to affect network properties (Zalesky et al., 2010). Because the resulting networks only capture dependencies between larger areas of cortex, this could obscure certain relationships only present at smaller scales. For example, we may have seen stronger effects in autism with a higher resolution atlas, in line with other studies showing a shift to short-range connectivity in this population (Weber et al., 2024). Again, the solution to this issue lies in larger sample sizes such that atlas downsampling is not necessary. It is also important to note that for some datasets, parcellation with the HCP atlas was also carried out using T1w scans only rather than the full HCP pipeline, which uses T2w and rs-fMRI scans. This did not appear to affect results, as datasets which underwent HCP pre-processing (e.g. PDC) did not differ in correlation structure from those that did not (e.g. DCAM). However, HCP preprocessing should ideally be applied across the board for methodological consistency, which will require open datasets to either provide preprocessed data or multimodal scans.

Finally, although this study makes an important contribution in applying the same methodological pipeline to various populations, it makes no direct comparison between these populations. This is due to both lowered statistical power as well as differences in age and sex balance between the populations. Empirical support for theories like the diametric model of autism and psychosis (Crespi and Badcock, 2008) will require direct comparison of high-level neurobiological features. Dealing with this issue will require larger datasets arising from collaboration between research groups, allowing pooling of subjects such that covariates like age and sex can be appropriately balanced.

## Acknowledgements

Data and/or research tools used in the preparation of this manuscript were obtained from the National Institute of Mental Health (NIMH) Data Archive (NDA). NDA is a collaborative informatics system created by the National Institutes of Health to provide a national resource to support and accelerate research in mental health. This manuscript reflects the views of the authors and may not reflect the opinions or views of the NIH or of the Submitters submitting original data to NDA. Data were provided (in part) by the Human Connectome Project, WU-Minn Consortium (Principal Investigators: David Van Essen and Kamil Ugurbil; 1U54MH091657) funded by the 16 NIH Institutes and Centers that support the NIH Blueprint for Neuroscience Research; and by the McDonnell Center for Systems Neuroscience at Washington University. Research reported in this publication was supported by the National Institute Of Mental Health of the National Institutes of Health under Award Number U01MH109589 and by funds provided by the McDonnell Center for Systems Neuroscience at Washington University in St. Louis. The HCP-Development 2.0 Release data used in this report came from DOI: 10.15154/1520708. Research reported in this publication was supported by the National Institute On Aging of the National Institutes of Health under Award Number U01AG052564 and by funds provided by the McDonnell Center for Systems Neuroscience at Washington University in St. Louis. The HCP-Aging 2.0 Release data used in this report came from DOI: 10.15154/1520707. Research using Human Connectome Project for Early Psychosis (HCP-EP) data reported in this publication was supported by the National Institute of Mental Health of the National Institutes of Health under Award Number U01MH109977. The HCP-EP 1.1 Release data used in this report came from DOI: 10.15154/1522899. Research reported in this publication was supported by the National Institute of Mental Health under Award Number U01MH108168 to Principal Investigators John D. E. Gabrieli and Susan Whitfield-Gabrieli. The BANDA 1.1 Release data used in this report came from DOI: 10.15154/3tk5-pb47. Research reported in this publication was supported by the National Institute of Mental Health of the National Institutes of Health under Award Number U01MH109991-01. The DCAM 1.0 Release data used in this report came from DOI: 10.15154/fj45-6d13. Research reported in this publication was supported by the National Institute of Mental Health of the National Institutes of Health under Award Number U01MH110008-04. The PDC 1.0 Release data used in this report came from DOI: 10.15154/1528673 For ABIDE-I, primary support for the work by Project Coordinator Adriana Di Martino was provided by the NIMH (K23MH087770) and the Leon Levy Foundation. Primary support for the work by Michael P. Milham and the INDI team was provided by gifts from Joseph P. Healy and the Stavros Niarchos Foundation to the Child Mind Institute, as well as by an NIMH award to MPM (NIMH R03MH096321). For ABIDE-II, primary support for the work by Project Coordinator Adriana Di Martino and her team was provided by the National Institute of Mental Health (NIMH 5R21MH107045). Primary support for the work by Michael P. Milham and his team provided by the National Institute of Mental Health (NIMH 5R21MH107045); Nathan S. Kline Institute of Psychiatric Research). Additional Support was provided by gifts from Joseph P. Healey, Phyllis Green and Randolph Cowen to the Child Mind Institute.

## Data and code availability

Analysis code and materials are available at https://github.com/wroseby/Global_SCNs_HC. The HCP-YA dataset is open access and available on the AWS S3 bucket hcp-openaccess/HCP_-1200/. Other HCP datasets are restricted access for researchers and are available upon application to the NDA. ABIDE-I and -II data are open access and are available on the AWS S3 bucket s3://fcp-indi/data/Projects/ABIDE/. The synaesthesia dataset is open-access and is available at https://www.nitrc.org/projects/syn_hcp/.

## Author contributions

WR contributed to data preprocessing, analysis, presentation, and manuscript writing. CR contributed to data preprocessing and analysis. JW conceived the study and provided feedback and supervision. All authors read and approved the final manuscript.

