## Supplementary Material for "Global organisation of structural covariance networks derived from parcellated cortical surface area in atypical populations"

| Sample | Clusters normal (%) | Clusters different variance (%) |
|---|---|---|
| Syn100 | Control: 85.48<br>Aff: 72.58 | 0.00 |
| ABIDE-I | Control: 74.19<br>Aff: 85.48 | 0.00 |
| ABIDE-II | Control: 70.97<br>Aff: 59.68 | 8.33 |
| HCP-EP | Control: 85.48<br>Aff: 79.03 | 1.11 |
| BANDA | Control: 79.03<br>Aff: 87.1 | 0.00 |
| DCAM | Control: 83.87<br>Aff: 61.29 | 2.78 |
| PDC | Control: 90.32<br>Aff: 79.03 | 0.00 |

**Table S1: Proportions of clusters passing tests for normality and group differences in variances.** Anderson-Darling test for normality and F-test for variance difference, $P < .05$ with FDR correction, $N = 102$.

| Cluster number | F-value | F-test P-value | Clustering | Global efficiency | Path length |
|---|---|---|---|---|---|
| 65 | 0.74 | <.001 | Control: 0.137<br>Syn: 0.158 | Control: 0.177<br>Syn: 0.193 | Control: 3.11<br>Syn: 2.87 |
| 64 | 0.76 | <.001 | Control: 0.137<br>Syn: 0.159 | Control: 0.177<br>Syn: 0.193 | Control: 3.10<br>Syn: 2.86 |
| 63 | 0.76 | <.001 | Control: 0.137<br>Syn: 0.158 | Control: 0.177<br>Syn: 0.195 | Control: 3.11<br>Syn: 2.84 |
| 61 | 0.81 | <.001 | Control: 0.138<br>Syn: 0.164 | Control: 0.178<br>Syn: 0.201 | Control: 3.09<br>Syn: 2.75 |
| 60 | 0.82 | <.001 | Control: 0.137<br>Syn: 0.164 | Control: 0.179<br>Syn: 0.201 | Control: 3.07<br>Syn: 2.76 |
| 59 | 0.83 | <.001 | Control: 0.135<br>Syn: 0.164 | Control: 0.177<br>Syn: 0.201 | Control: 3.11<br>Syn: 2.75 |

**Table S2: Group differences in partial correlation variance and network metrics across different numbers of clusters in the Syn100 dataset.** Different numbers of clusters were selected by varying the threshold for cutting the hierarhical clustering dendrogram around the optimal.

| Sample | Clustering | Clustering effect size | Clustering P-value | Global efficiency | Global efficiency effect size | Global efficiency P-value | Path length | Path length effect size | Path length P-value |
|---|---|---|---|---|---|---|---|---|---|
| Syn100 | Control: 0.037<br>Aff: 0.038 | 0.16 | .4299 | Control: 0.156<br>Aff: 0.172 | 1.53 | .0631 | Control: 3.539<br>Aff: 3.205 | -1.58 | .0605 |
| ABIDE-I | Control: 0.038<br>Aff: 0.034 | -1.34 | .9183 | Control: 0.170<br>Aff: 0.182 | 1.13 | .1274 | Control: 3.271<br>Aff: 3.062 | -1.13 | .1282 |
| ABIDE-II | Control: 0.037<br>Aff: 0.036 | -0.43 | .6767 | Control: 0.168<br>Aff: 0.183 | 1.41 | .0771 | Control: 3.323<br>Aff: 3.042 | -1.56 | .0592 |
| HCP-EP | Control: 0.037<br>Aff: 0.036 | -0.33 | .3648 | Control: 0.178<br>Aff: 0.171 | -0.62 | .2611 | Control: 3.115<br>Aff: 3.254 | 0.76 | .2215 |
| BANDA | Control: 0.038<br>Aff: 0.040 | 0.73 | .2205 | Control: 0.158<br>Aff: 0.161 | 0.27 | .3918 | Control: 3.516<br>Aff: 3.434 | -0.4 | .3455 |
| DCAM | Control: 0.036<br>Aff: 0.037 | 0.18 | .5658 | Control: 0.167<br>Aff: 0.170 | 0.33 | .6332 | Control: 3.320<br>Aff: 3.266 | -0.27 | .6067 |
| PDC | Control: 0.038<br>Aff: 0.040 | 0.63 | .7445 | Control: 0.171<br>Aff: 0.171 | 0.04 | .5129 | Control: 3.235<br>Aff: 3.225 | -0.04 | .5155 |

**Table S3: Group differences in global network metrics for positive-only SCNs.** P-values were calculated via permutation tests with $n = 10,000$ permutations. Effect size for each measure is calculated using the observed group difference in metrics normalised by the null (shuffled) distribution standard deviation.

| Sample | Clustering | Clustering effect size | Clustering P-value | Global efficiency | Global efficiency effect size | Global efficiency P-value | Path length | Path length effect size | Path length P-value |
|---|---|---|---|---|---|---|---|---|---|
| Syn100 | Control: 0.092<br>Aff: 0.115 | 1.32 | .0838 | Control: 0.122<br>Aff: 0.138 | 1.83 | **.0336** | Control: 4.680<br>Aff: 4.178 | -1.85 | **.0328** |
| ABIDE-I | Control: 0.087<br>Aff: 0.108 | 1.13 | .1147 | Control: 0.126<br>Aff: 0.138 | 1.24 | .1057 | Control: 4.475<br>Aff: 4.153 | -1.28 | .0995 |
| ABIDE-II | Control: 0.089<br>Aff: 0.110 | 1.1 | .1188 | Control: 0.127<br>Aff: 0.138 | 1.2 | .1099 | Control: 4.421<br>Aff: 4.133 | -1.17 | .1189 |
| HCP-EP | Control: 0.111<br>Aff: 0.101 | -0.53 | .2786 | Control: 0.138<br>Aff: 0.129 | -1.03 | .1503 | Control: 4.127<br>Aff: 4.426 | 1.24 | .1088 |
| BANDA | Control: 0.096<br>Aff: 0.096 | -0.03 | .5179 | Control: 0.125<br>Aff: 0.129 | 0.43 | .3341 | Control: 4.559<br>Aff: 4.413 | -0.56 | .2918 |
| DCAM | Control: 0.099<br>Aff: 0.102 | 0.13 | .5640 | Control: 0.129<br>Aff: 0.127 | -0.2 | .4215 | Control: 4.412<br>Aff: 4.522 | 0.47 | .3184 |
| PDC | Control: 0.109<br>Aff: 0.112 | 0.16 | .5695 | Control: 0.135<br>Aff: 0.137 | 0.34 | .6394 | Control: 4.244<br>Aff: 4.180 | -0.24 | .5951 |

**Table S4: Group differences in global network metrics for negative-only SCNs.** P-values were calculated via permutation tests with $n = 10,000$ permutations. Effect size for each measure is calculated using the observed group difference in metrics normalised by the null (shuffled) distribution standard deviation.

| **Sample** | **Δ mean** | **Δ mean P-value** | **Δ variance** | **Δ variance P-value** | **Δ skew** | **Δ skew P-value** | **Δ kurtosis** | **Δ kurtosis P-value** |
|---|---|---|---|---|---|---|---|---|
| Syn100 | -0.001 | .6299 | 0.000 | .0652 | 1.063 | .0825 | 3.317 | .1914 |
| ABIDE-I | 0.000 | .5195 | 0.000 | .3342 | 0.167 | .3996 | 0.151 | .4853 |
| ABIDE-II | -0.001 | .6410 | 0.000 | .2069 | -0.061 | .5740 | -0.685 | .7858 |
| HCP-EP | 0.001 | .2646 | 0.000 | .5192 | -0.334 | .8415 | -0.771 | .7579 |
| BANDA | -0.001 | .6802 | 0.000 | .5319 | -0.059 | .5593 | -0.396 | .6361 |
| DCAM | 0.001 | .2364 | 0.000 | .0838 | 0.542 | .0510 | 1.005 | .1540 |
| PDC | 0.000 | .4445 | 0.000 | .4128 | 0.271 | .2317 | 1.192 | .1583 |

**Table S5: Group differences in betweenness centrality distributions for positive-only SCNs.** P-values were calculated via permutation tests with $n = 10,000$ permutations; $n = 62$ observations for all tests.

| **Sample** | **Δ mean** | **Δ mean P-value** | **Δ variance** | **Δ variance P-value** | **Δ skew** | **Δ skew P-value** | **Δ kurtosis** | **Δ kurtosis P-value** |
|---|---|---|---|---|---|---|---|---|
| Syn100 | 0.001 | .2850 | 0.000 | **.0316** | 0.673 | .3195 | 2.560 | .3951 |
| ABIDE-I | 0.001 | .4029 | 0.000 | .4606 | -0.196 | .7202 | 0.061 | .4641 |
| ABIDE-II | 0.001 | .3901 | 0.000 | .1160 | -0.214 | .6565 | -0.768 | .6598 |
| HCP-EP | -0.001 | .5839 | 0.000 | .9803 | -1.186 | .9301 | -3.681 | .7358 |
| BANDA | 0.000 | .4554 | 0.000 | .7184 | -0.475 | .7501 | -1.881 | .7226 |
| DCAM | 0.001 | .4296 | 0.000 | .1558 | 0.240 | .3207 | -0.552 | .6088 |
| PDC | 0.002 | .2565 | 0.000 | .2186 | 0.340 | .2867 | 1.772 | .2665 |

**Table S6: Group differences in betweenness centrality distributions for negative-only SCNs.** P-values were calculated via permutation tests with $n = 10,000$ permutations; $n = 62$ observations for all tests.

| Sample | FC network F(6, 3775) | FC network P-value | Visual t | Visual P-value | Somatomotor t | Somatomotor P-value | DA t | DA P-value | VA t | VA P-value | Limbic t | Limbic P-value | FP t | FP P-value | Default t | Default P-value |
|---|---|---|---|---|---|---|---|---|---|---|---|---|---|---|---|---|
| Syn100 | 3.97 | **.0006** | -0.08 | .9390 | 0.09 | .9283 | -0.12 | .9083 | 0.51 | .9116 | 0.51 | .6110 | 0.15 | .8811 | -0.27 | .7855 |
| ABIDE-I | 0.46 | .8379 | 0.12 | .9018 | -0.03 | .9777 | 0.01 | .9937 | -0.23 | .8090 | -0.23 | .8192 | 0.18 | .8566 | -0.30 | .7630 |
| ABIDE-II | 2.96 | **.0070** | -0.05 | .9589 | -0.21 | .8341 | -0.01 | .9926 | -0.11 | .7942 | -0.11 | .9086 | -0.12 | .9075 | 0.19 | .8530 |
| HCP-EP | 3.19 | **.0040** | 0.19 | .8478 | 0.13 | .8974 | 0.14 | .8894 | 0.13 | .9064 | 0.13 | .9002 | -0.22 | .8275 | -0.18 | .8547 |
| BANDA | 2.67 | **.0137** | -0.05 | .9618 | -0.40 | .6889 | -0.08 | .9330 | 0.01 | .6950 | 0.01 | .9903 | 0.10 | .9170 | 0.04 | .9667 |
| DCAM | 5.18 | $<$ **.0001** | -0.05 | .9635 | -0.09 | .9257 | 0.05 | .9583 | -0.16 | .9084 | -0.16 | .8709 | 0.03 | .9728 | 0.05 | .9636 |
| PDC | 1.44 | .1952 | -0.26 | .7970 | 0.20 | .8423 | -0.30 | .7655 | 0.33 | .9666 | 0.33 | .7444 | -0.31 | .7600 | 0.25 | .8009 |

**Table S7: Comparison of network weights across seven functional connectivity networks.** Clusters were assigned to a network according to which network the majority of their parcels belonged. Each weight was assigned to two networks based on the consensus membership of its clusters. Mean group differences in weight (Z-scores) were compared across networks via one-way ANOVA, then the mean of each FC network was compared against that of all others via t-test. FC = functional connectivity, DA = dorsal attention, VA = ventral attention, FP = frontoparietal.

| Sample | Group coefficient | Group coefficient SE | Group coefficient $t_{(3772.363)}$ | Group coefficient P-value | Smooth term F | Smooth term P-value | Group:smooth deviance | Group:smooth P-value | n obs |
|---|---|---|---|---|---|---|---|---|---|
| Syn100 | 0.014 | 0.005 | 3.074 | **.0021** | Control: 30.64<br>Aff: 30.06 | Control: $<$ **.0001**<br>Aff: $<$ **.0001** | 0.093 | .0647 | Control: 1009<br>Aff: 1014 |
| ABIDE-I | 0.006 | 0.005 | 1.268 | .2051 | Control: 74.02<br>Aff: 85.14 | Control: $<$ **.0001**<br>Aff: $<$ **.0001** | 0.051 | .3616 | Control: 958<br>Aff: 1005 |
| ABIDE-II | 0.012 | 0.005 | 2.507 | **.0122** | Control: 92.03<br>Aff: 72.52 | Control: $<$ **.0001**<br>Aff: $<$ **.0001** | 0.055 | .3170 | Control: 985<br>Aff: 987 |
| HCP-EP | -0.012 | 0.005 | -2.457 | **.0141** | Control: 57.86<br>Aff: 64.86 | Control: $<$ **.0001**<br>Aff: $<$ **.0001** | 0.035 | .5593 | Control: 986<br>Aff: 995 |
| BANDA | 0.007 | 0.004 | 1.606 | .1085 | Control: 26.91<br>Aff: 21.13 | Control: $<$ **.0001**<br>Aff: $<$ **.0001** | 0.040 | .4001 | Control: 1024<br>Aff: 997 |
| DCAM | 0.002 | 0.005 | 0.411 | .6814 | Control: 44.81<br>Aff: 59.08 | Control: $<$ **.0001**<br>Aff: $<$ **.0001** | 0.018 | .8004 | Control: 999<br>Aff: 986 |
| PDC | -0.003 | 0.005 | -0.604 | .5461 | Control: 19.97<br>Aff: 14.06 | Control: $<$ **.0001**<br>Aff: $<$ **.0001** | 0.028 | .6264 | Control: 980<br>Aff: 1003 |

**Table S8: Generalised additive models (GAM) predicting positive network weights as a function of a parametric Group term, and separate smooth Distance terms for each group.** Estimated and reference degrees of freedom for F-values can be found in Table S9.

| Sample | Group coefficient | Group coefficient SE | Group coefficient $t_{(3772.363)}$ | Group coefficient P-value | Smooth term F | Smooth term P-value | Group:smooth deviance | Group:smooth P-value | n obs |
|---|---|---|---|---|---|---|---|---|---|
| Syn100 | -0.018 | 0.005 | -4.013 | $<$ **.0001** | Control: 0.38<br>Aff: 0.59 | Control: .5357<br>Aff: .4434 | 0.009 | .3268 | Control: 882<br>Aff: 877 |
| ABIDE-I | -0.020 | 0.004 | -4.411 | $<$ **.0001** | Control: 3.70<br>Aff: 8.60 | Control: **.0192**<br>Aff: **.0034** | 0.016 | .2323 | Control: 933<br>Aff: 886 |
| ABIDE-II | -0.015 | 0.004 | -3.361 | **.0008** | Control: 3.77<br>Aff: 3.21 | Control: **.0191**<br>Aff: **.0373** | -0.007 | nan | Control: 906<br>Aff: 904 |
| HCP-EP | 0.011 | 0.005 | 2.301 | **.0215** | Control: 4.15<br>Aff: 2.63 | Control: **.0062**<br>Aff: **.0299** | -0.006 | nan | Control: 905<br>Aff: 896 |
| BANDA | 0.000 | 0.004 | -0.038 | .9698 | Control: 0.90<br>Aff: 0.43 | Control: .3428<br>Aff: .5144 | 0.000 | .8304 | Control: 867<br>Aff: 894 |
| DCAM | 0.001 | 0.004 | 0.287 | .7741 | Control: 0.34<br>Aff: 1.68 | Control: .5619<br>Aff: .1236 | 0.044 | .0691 | Control: 892<br>Aff: 905 |
| PDC | -0.004 | 0.005 | -0.912 | .3621 | Control: 2.62<br>Aff: 1.31 | Control: .1060<br>Aff: .2000 | 0.026 | .1445 | Control: 911<br>Aff: 888 |

**Table S9: Generalised additive models (GAM) predicting negative network weights as a function of a parametric Group term, a smooth Distance term and a Group:smooth Distance interaction.** Estimated and reference degrees of freedom for F values can be found in Table S9.

| Sample | F edf | F ref | F edf (pos) | F ref (pos) | F edf (neg) | F ref (neg) |
|---|---|---|---|---|---|---|
| Syn100 | Control: 3.72<br>Aff: 3.92 | Control: 3.96<br>Aff: 3.96 | Control: 3.73<br>Aff: 3.93 | Control: 3.96<br>Aff: 4.00 | Control: 1.00<br>Aff: 1.00 | Control: 1.00<br>Aff: 1.00 |
| ABIDE-I | Control: 3.94<br>Aff: 3.95 | Control: 4.00<br>Aff: 4.00 | Control: 3.92<br>Aff: 3.92 | Control: 4.00<br>Aff: 4.00 | Control: 1.87<br>Aff: 1.00 | Control: 2.32<br>Aff: 1.00 |
| ABIDE-II | Control: 3.92<br>Aff: 3.94 | Control: 4.00<br>Aff: 4.00 | Control: 3.88<br>Aff: 3.93 | Control: 4.00<br>Aff: 4.00 | Control: 1.95<br>Aff: 1.69 | Control: 2.42<br>Aff: 2.09 |
| HCP-EP | Control: 3.93<br>Aff: 3.91 | Control: 4.00<br>Aff: 4.00 | Control: 3.94<br>Aff: 3.91 | Control: 4.00<br>Aff: 4.00 | Control: 2.48<br>Aff: 3.27 | Control: 3.00<br>Aff: 3.72 |
| BANDA | Control: 3.79<br>Aff: 3.85 | Control: 3.97<br>Aff: 3.99 | Control: 3.78<br>Aff: 3.68 | Control: 3.97<br>Aff: 3.94 | Control: 1.00<br>Aff: 1.00 | Control: 1.00<br>Aff: 1.00 |
| DCAM | Control: 3.91<br>Aff: 3.94 | Control: 4.00<br>Aff: 4.00 | Control: 3.87<br>Aff: 3.91 | Control: 4.00<br>Aff: 3.91 | Control: 1.00<br>Aff: 2.35 | Control: 1.00<br>Aff: 2.86 |
| PDC | Control: 3.84<br>Aff: 3.57 | Control: 3.98<br>Aff: 3.89 | Control: 3.81<br>Aff: 3.41 | Control: 3.98<br>Aff: 3.81 | Control: 1.00<br>Aff: 2.99 | Control: 1.00<br>Aff: 3.50 |

**Table S10: Degrees of freedom for F-tests across all GAMs.** F-tests were performed for the significance of the smooth term of weight as a function of distance. ‘edf’ = estimated degrees of freedom, ‘ref’ = reference degrees of freedom.

| Sample | SWP (positive) | delta (positive) | SWP (negative) | delta (negative) |
|---|---|---|---|---|
| Syn100 | Control: 0.71<br>Aff: 0.71 | Control: -1.00<br>Aff: -0.99 | Control: 0.38<br>Aff: 0.32 | Control: -0.51<br>Aff: -0.26 |
| ABIDE-I | Control: 0.71<br>Aff: 0.71 | Control: -1.00<br>Aff: -0.99 | Control: 0.42<br>Aff: 0.35 | Control: -0.73<br>Aff: -0.43 |
| ABIDE-II | Control: 0.71<br>Aff: 0.71 | Control: -0.98<br>Aff: -0.99 | Control: 0.41<br>Aff: 0.36 | Control: -0.70<br>Aff: -0.39 |
| HCP-EP | Control: 0.71<br>Aff: 0.71 | Control: -0.97<br>Aff: -0.98 | Control: 0.33<br>Aff: 0.36 | Control: -0.39<br>Aff: -0.46 |
| BANDA | Control: 0.71<br>Aff: 0.71 | Control: -0.98<br>Aff: -1.00 | Control: 0.37<br>Aff: 0.38 | Control: -0.48<br>Aff: -0.57 |
| DCAM | Control: 0.71<br>Aff: 0.71 | Control: -0.99<br>Aff: -1.00 | Control: 0.34<br>Aff: 0.35 | Control: -0.57<br>Aff: -0.35 |
| PDC | Control: 0.71<br>Aff: 0.71 | Control: -0.97<br>Aff: -1.00 | Control: 0.33<br>Aff: 0.33 | Control: -0.37<br>Aff: -0.35 |

**Table S11:** Small-world propensity (SWP) and delta values for positive- and negative-only SCNs.

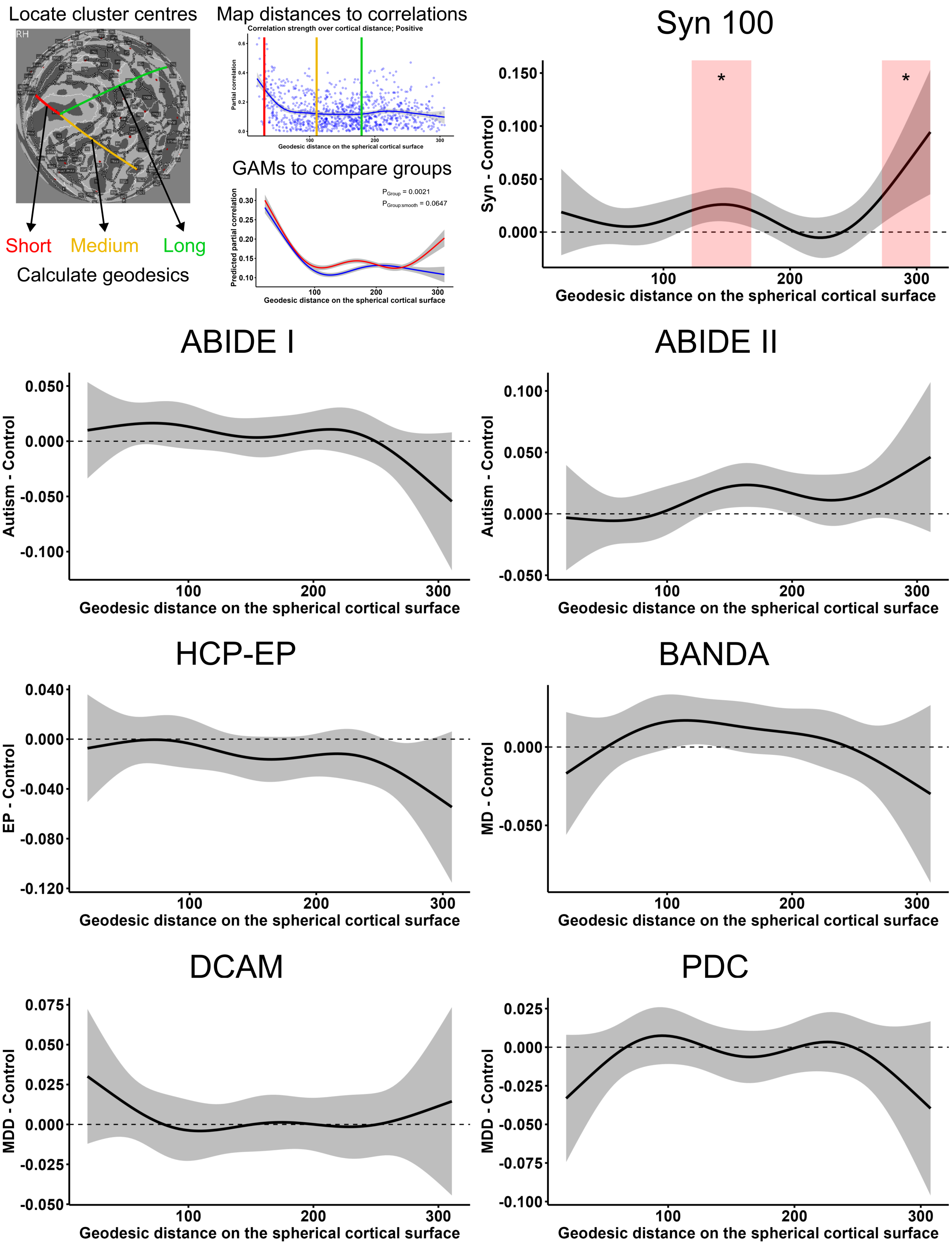


**Figure S1: Group differences in positive-only network weights over the range of inter-cluster cortical distances.** GAMs were used used to model the smooth relationship between cortical distance and network weight, according to group. The plots for each dataset show the group difference in predicted network weights across the range of cortical distances, with the shaded area indicating the $95\%$ confidence interval. Distances with significant group differences are highlighted; '*' = $P < .05$, with FDR correction for 100 tests.

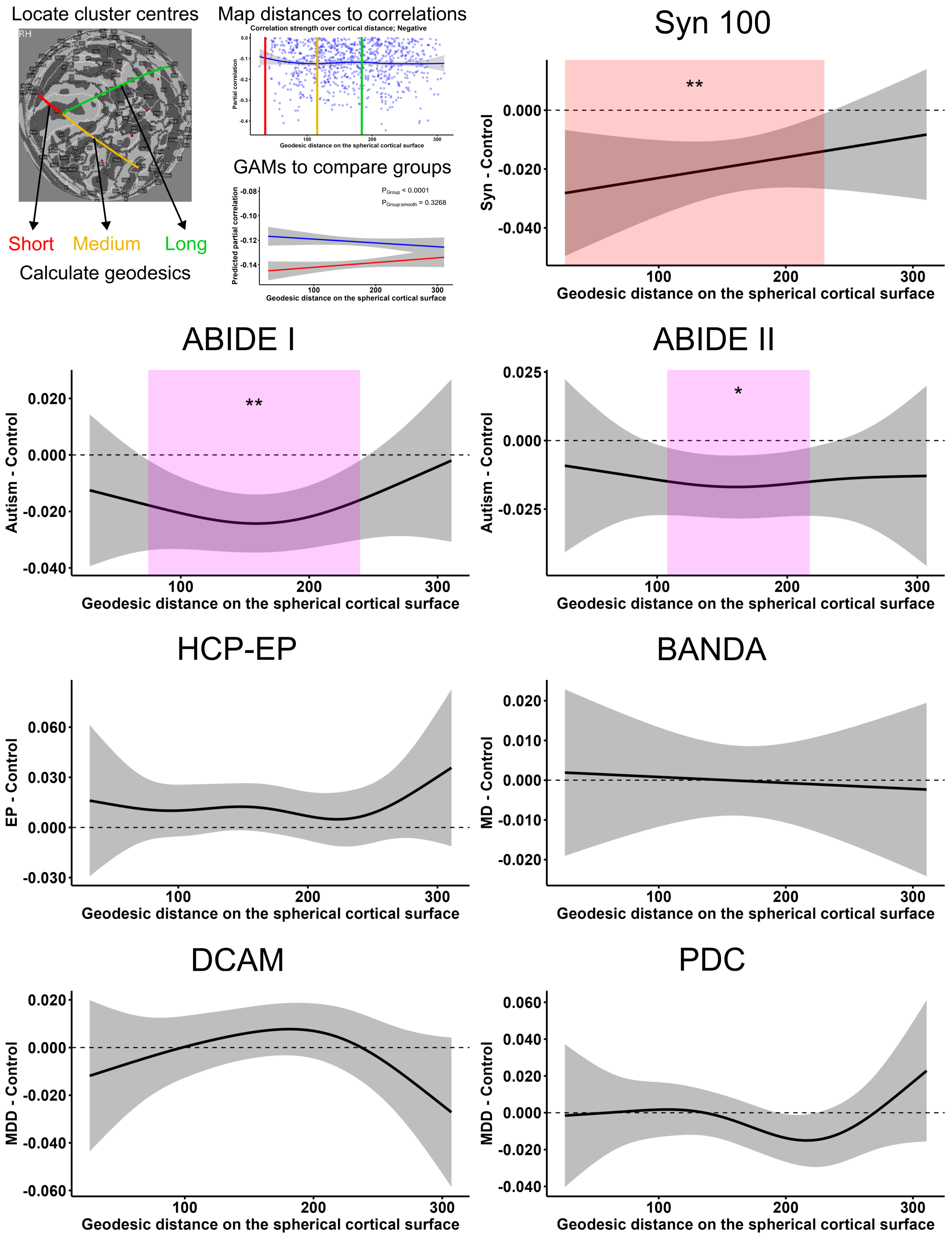


**Figure S2: Group differences in negative-only network weights over the range of inter-cluster cortical distances.** GAMs were used used to model the smooth relationship between cortical distance and network weight, according to group. The plots for each dataset show the group difference in predicted network weights across the range of cortical distances, with the shaded area indicating the $95\%$ confidence interval. Distances with significant group differences are highlighted; '**' = $P < .01$, '*' = $P < .05$, with FDR correction for 100 tests. GAMs were fitted with $n = 1891$ observations per group.
